# Functional regulation of an intrinsically disordered protein via a conformationally excited state

**DOI:** 10.1101/2022.06.24.497365

**Authors:** Kulkarni Madhurima, Bodhisatwa Nandi, Sneha Munshi, Athi N. Naganathan, Ashok Sekhar

**Author notes:** Present address: Department of Physics, University of Alberta, Edmonton, AB T6G 2E1, Canada.

## Abstract

A longstanding goal in the field of intrinsically disordered proteins (IDP) is to characterize their structural heterogeneity and pinpoint the role of this heterogeneity in IDP function. Here, we use multinuclear chemical exchange saturation (CEST) NMR to determine the structure of a thermally accessible globally folded excited state in equilibrium with the intrinsically disordered native ensemble of a bacterial transcriptional regulator CytR. We further provide evidence from double resonance CEST experiments that the excited state, which structurally resembles the DNA-bound form of CytR, recognizes DNA by means of a ‘folding-before-binding’ conformational selection pathway. The disorder-to-order regulatory switch in DNA recognition by natively disordered CytR therefore operates through a dynamical variant of the lock-and-key mechanism where the structurally complementary conformation is transiently accessed via thermal fluctuations.

**One-Sentence Summary:** The intrinsically disordered cytidine repressor binds DNA via a folding-before-binding conformational selection mechanism

---

Between 16-45 % of bacterial and over 50 % of eukaryotic proteins contain long intrinsically disordered protein regions (IDPs), which do not possess stable structure in isolation but instead exist as a heterogeneous ensemble of interconverting conformations (*1*). The structural plasticity of IDPs makes them particularly suited for participating in signal transduction cascades and regulatory networks, and disorder is prevalently found in kinases, transcription factors and nucleic acid-binding proteins (*2, 3*). The primary sequences of IDPs are simpler than sequences of folded proteins and are characterized by low complexity coupled with a biased amino acid composition (*1*). While small (Gly, Ser) and charged amino acids (Lys, Arg, Asp, Glu) are enriched in IDPs, there is a simultaneous depletion of large aliphatic (Leu, Ile and Val) and aromatic (Phe, Tyr, Trp) hydrophobic residues which promote order and folding (*3*). The conformational tendencies of an IDP are largely dictated by its amino acid sequence and the conformational free energy surfaces of IDPs are believed to be flat and lacking pronounced minima (*4–6*).

Despite being unable to fold spontaneously into stable three-dimensional structures, IDPs have evolved distinctive mechanisms for performing their function. IDPs frequently undergo disorder-to-order transitions that result from interactions with binding partners(*7, 8*) or as a consequence of post-translational modifications(*9*). In addition, disordered regions have been shown to nucleate liquid-liquid phase separation(*10–12*), thereby creating dynamic membrane-less organelles for compartmentalizing the localization and function of cellular components. The fundamental motif underlying IDP function is biomolecular recognition, which is enabled by the weak, multivalent and often promiscuous interactions of IDPs with themselves and with other physiological binding partners(*12, 13*).

A central challenge in IDP biophysics has been the difficulty in obtaining an atomic-resolution description of the mechanisms of interaction between IDPs and their partner proteins. The widely accepted mechanism of recognition is the folding-upon-binding pathway(*14, 15*), where transient encounter complexes stabilized by native or non-native interactions between IDPs and their partners mature into the final protein-protein or protein-nucleic acid complex without dissociation of the IDP from its binding partner (*16–18*). This pathway requires no prior folding of the IDP (*19*)and while IDPs have been known to possess small amounts of pre-formed secondary structure(*20*), such residual structure is not required for binding (*21*) and often has to unfold before the binding event (*17*).

In this report, we show using multinuclear chemical exchange saturation transfer (CEST) NMR that the natively disordered N-terminal domain of the cytidine repressor, belonging to the LacR family of transcriptional regulators, transiently populates a globally folded conformationally excited state. We use chemical shifts and residual dipolar couplings to determine the structure of the excited state, which is a well-organized three-helix bundle containing a helix-turn-helix DNA recognition motif. We then employ the recently developed multifrequency irradiation Double Resonance DANTE-CEST to demonstrate a novel mode of IDP functional regulation, in which the excited state alone binds DNA through a folding-before-binding conformational selection mechanism.

## Results

### CytR^N^ is an intrinsically disordered DNA binding domain

The cytidine repressor (CytR) (*22*) is a member of the LacR family of transcriptional repressors(*23, 24*). It is 341 amino acids long and consists of a 66-residue N-terminal DNA binding domain (DBD, referred to as CytR^N^), followed by a C-terminal region responsible for binding cytidine as well as dimerization(*25*). While the DNA binding domains of members of the LacR family including LacR, FruR, and PurR fold into stable three-dimensional structures in the absence of DNA (*26–28*), sequence-specific disorder prediction algorithms PONDR(*29, 30*) and IUPRED(*31*) classify CytR^N^ as an intrinsically disordered protein region (Fig. S1A). CytR^N^ has significantly higher mean net charge and lower hydrophobicity than its family members that places it in the disordered region of the Uversky charge-hydropathy plot (Fig. 1A). In addition, CytR^N^ is depleted in hydrophobic residues that promote ordering and is instead rich in Ala and Lys which are abundant in disordered proteins (Fig. S1B). Figure 1B shows the ^1^H-^15^N HSQC spectrum of CytR^N^ in its native state. The backbone amide resonances in the ^1^H dimension are dispersed over a narrow chemical shift range between 7.9 and 8.8 ppm, confirming that CytR^N^ is intrinsically disordered. Residue-specific secondary structure propensity (SSP(*32*)) scores, calculated using backbone chemical shifts, indicate that native CytR^N^ does not adopt stable secondary structure, though it has up to ∼30 % residual helicity in regions of the protein that form helices in the DNA-bound conformation (Fig. 1C) (*33*).

**Fig. 1.**
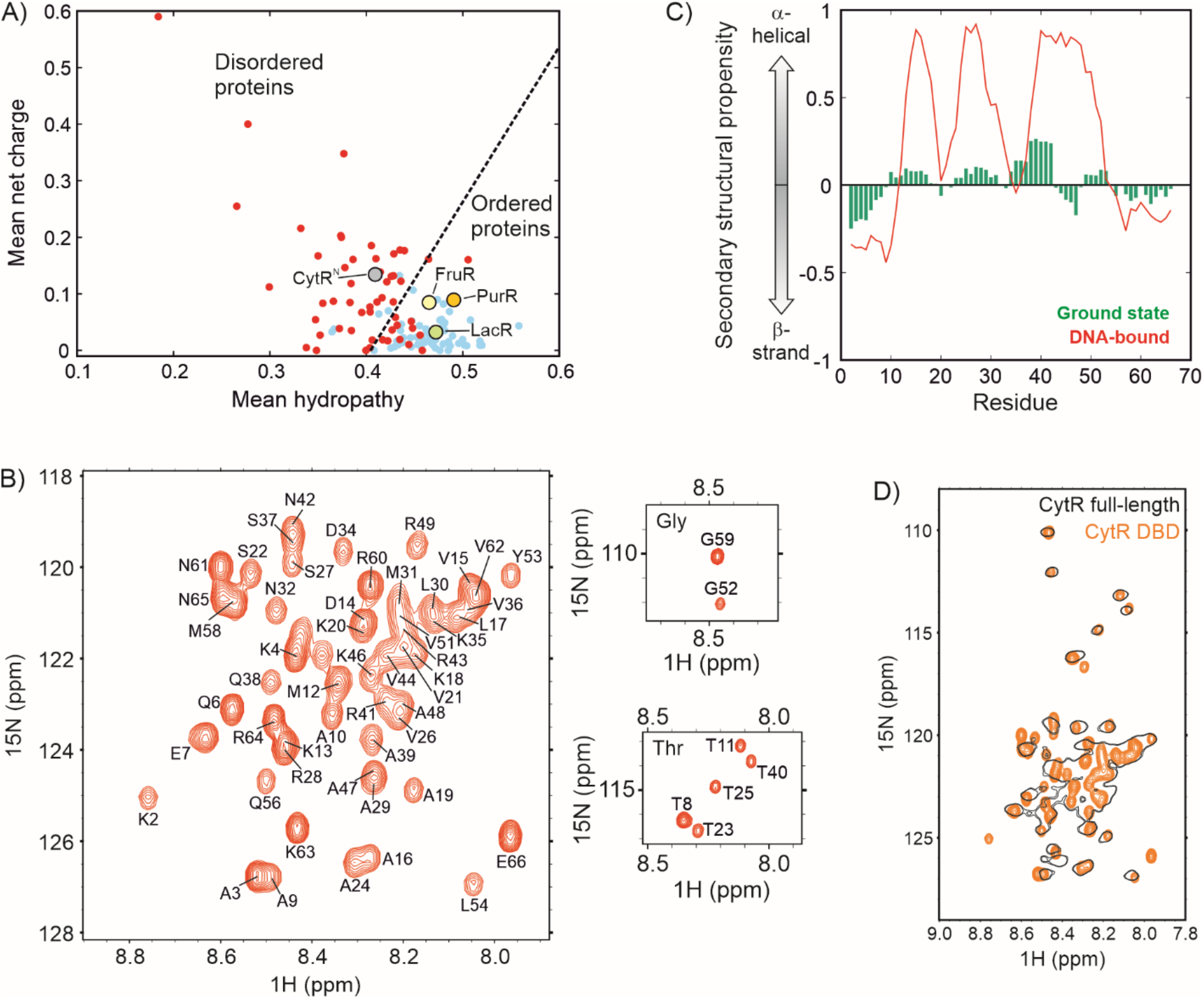
CytR^N^ is intrinsically disordered by itself and in the full-length cytidine repressor. A) Charge versus hydropathy scores show that CytR^N^ falls in the region occupied by disordered proteins (red dots), while LacR, FruR and PurR fall in the region of ordered proteins (blue dots). B) The ^1^H-^15^N HSQC spectrum of CytR^N^. The resonance assignments for each peak are indicated on the spectrum. C) Secondary structure propensity of native disordered (green) and DNA-bound CytR^N^ (red). D) Overlay of ^1^H-^15^N HSQC spectra of CytR^N^ (orange) and full-length CytR (black).

In order to determine whether native CytR^N^ is unable to fold because of unsatisfied interactions with the C-terminal domain that are absent in the truncated version, we acquired a ^1^H-^15^N HSQC spectrum of full-length CytR which exists as a dimer of 38 kDa protomers. Most of the resonances visible in the HSQC spectrum of CytR (Fig. 1D, black contours) overlay very well with matching resonances from CytR^N^ (Fig. 1D, orange), indicating that the N-terminal DBD is unstructured in full-length CytR also. Although resonances from structured regions of the 76 kDa CytR are expected to be broadened out in the HSQC spectrum because of slow tumbling and large transverse relaxation rate constants, peaks from the N-terminus are sharp, demonstrating that CytR^N^ behaves independently of the rest of the protein and retains considerable local mobility in full-length CytR. Taken together, NMR data confirm predictions based on the sequence of CytR^N^ that native CytR^N^ is intrinsically disordered in both the full-length and truncated versions of the protein.

### CytR^N^ transiently populates a higher free energy excited state

Proteins are dynamic and often utilize thermally accessible higher free energy conformations (referred to here as excited states) for performing function(*34*). However, our current understanding suggests that excited conformations of IDPs are markedly destabilized with respect to the native disordered ensemble, because the latter exists as a heterogeneous collection of microstates with comparable energies that increases its entropy(*4*). Therefore, the populations of excited states of IDPs are expected to be too low to be detectable or functional. Nevertheless, previous fluorescence-detected stopped-flow experiments provided evidence that CytR^N^ may be undergoing compaction on the millisecond timescale(*35*). Hence, we next probed the slow dynamics of CytR^N^ using Chemical Exchange Saturation Transfer (CEST) (*36*) NMR.

CEST experiments were carried out by irradiating ^15^N-labeled CytR^N^ with a weak radiofrequency field of amplitude B_1_ over a range of ^15^N offsets for a fixed exchange duration (T_ex_), “searching” for chemical shifts of NMR spins in invisible excited states. These can be ascertained by quantifying the intensities (I) of peaks from the disordered state in CEST profiles, which are graphs of the ratio of I with the corresponding intensity in a reference experiment where the exchange duration is not present. The fingerprint of conformational exchange in CEST profiles is the appearance of two or more dips, the larger one arising from saturation of the major state and the smaller one, from perturbations of the minor state by the B_1_ field which are subsequently amplified and transferred to the major state. Since the minor dip in intensity occurs at the chemical shift of a spin in the excited state, CEST profiles provide an avenue for determining the chemical shifts of ‘invisible’ conformations(*36*) that cannot be detected in routine NMR spectra.

The presence of a kinetically distinct excited state in the thermal ensemble of CytR^N^ can be discerned from ^15^N CEST profiles of a number of residues (Fig. 2A, Fig. S2) as two dips in intensity. CEST data acquired at two B_1_ fields (14.7 and 28.6 Hz) for 12 isolated residues showing two well-separated dips were globally modeled using the Bloch-McConnell equations. Fits of the CEST data reveal that the population of the excited state (E) is 8.7 ± 0.1 % at 288 K (Fig. S3), which places this conformation at 1.34 kcal/mol (or 2.35 kT) higher in free energy than the native disordered form (D, Fig. 2B). The exchange rate constant between the two states (k_ex,DE_) is 45.9 ± 0.9 s^-1^, which implies that the excited state is transiently populated with a lifetime of 23.8 ± 0.5 ms (Fig. S3). Our data thus unequivocally demonstrate that, contrary to the currently accepted notions, IDPs can access excited conformations with millisecond lifetimes via thermal fluctuations.

**Fig. 2.**
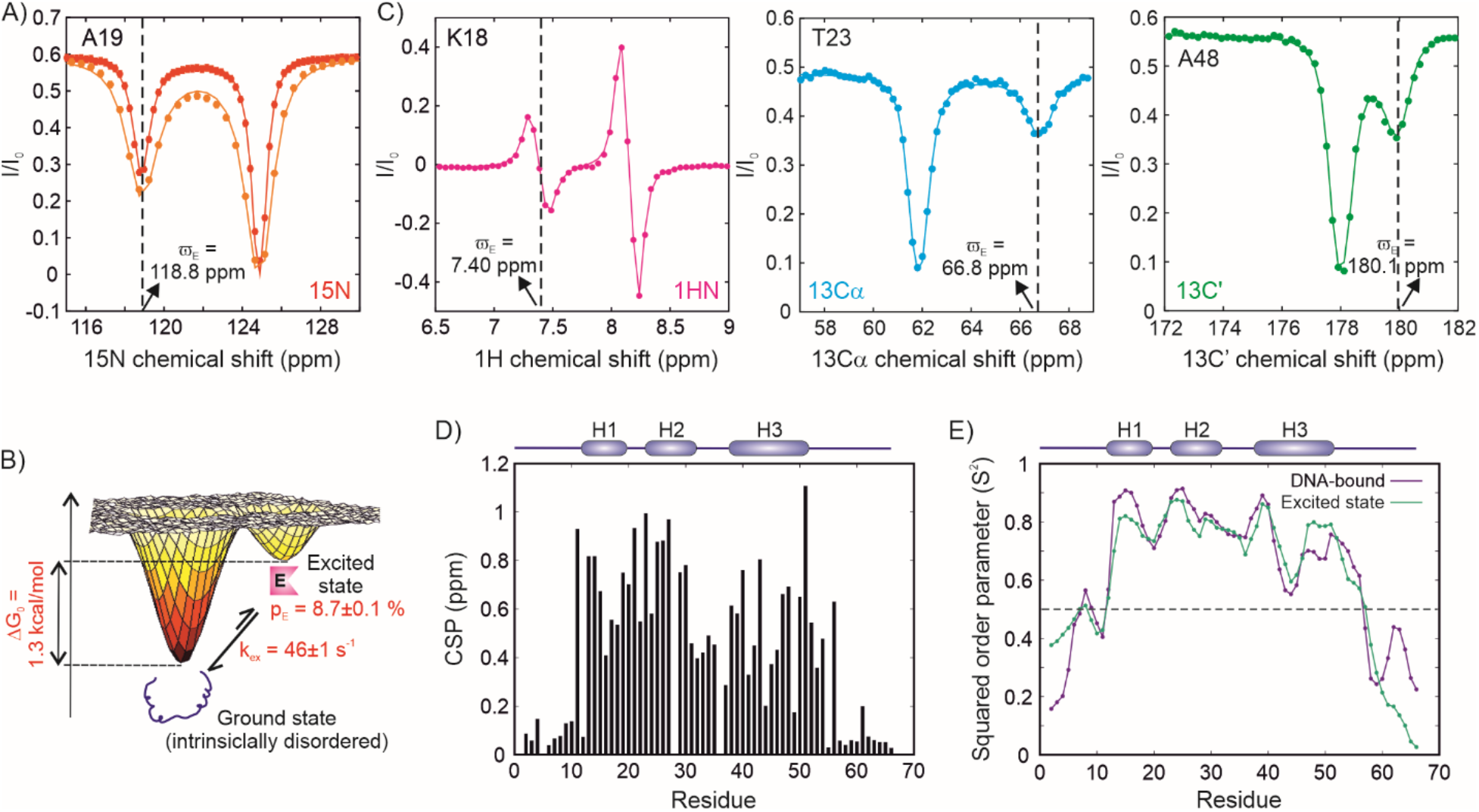
Intrinsically disordered CytR^N^ populates a globally folded excited state. A) ^15^N CEST profiles of A19, acquired at 14.7 Hz (red) and 28.6 Hz (orange). Solid lines are global fits of the CEST data to the Bloch-McConnell equations. The dashed line shows the chemical shift position of the nucleus in the excited state. B) Cartoon representation of the conformational free energy surface of CytR^N^, showing the intrinsically disordered ground state in equilibrium with an excited state. C) CEST profiles of the amide proton of K18 (left, magenta), alpha carbon of T23 (middle, cyan) and carbonyl carbon of A48 (right, green). The solid lines are fits to the Bloch-McConnell equations while the dashed lines indicate the chemical shift position of the excited state dip. D) Chemical shift perturbations (CSP) between the excited and ground states of CytR^N^, calculated as described in the Materials and Methods. E) Residue-specific squared order parameters (S^2^) for the excited state (green) and DNA-bound CytR^N^ (purple). The secondary structural elements of DNA-bound CytR^N^ are shown at the top of the plot in panels D and E.

### The excited state of CytR^N^ is globally folded

IDPs have been known to undergo local folding as well as global disorder-to-order transitions in the presence of binding partners or upon post-translational modifications. In order to determine the extent of structure in the excited state of CytR^N^, we used multinuclear CEST profiles to extract ^15^N(*37*), ^1^H^N^ (*38*), ^13^Cα (*39*) and ^13^CO(*40*) chemical shifts in the excited conformation (Fig. 2C, Fig. S4). ^15^N, ^1^H^N, 13^Cα and ^13^CO chemical shifts were obtained for 53, 51, 40 and 44 residues respectively (Tables S2 and S3), providing excellent coverage of the structural changes occurring in the entirety of CytR^N^.

Since chemical shifts are extremely sensitive probes of protein structure(*41*), regions of the protein experiencing large chemical shift perturbations (CSPs) will also be the regions that display large differences in conformation between the excited state and the native disordered ensemble. Figure 2D shows the residue-specific CSPs in CytR^N^ resulting from the conformational interconversion. CSPs are small (< 0.2 ppm) at the N-(M1-A10) and C-termini (P57-E66) and signify that the extremities of CytR^N^ remain disordered in the excited state. Interestingly, while residues 50-57 in LacR form the hinge helix when the repressor is bound to operator DNA(*42*), the corresponding region in CytR remains disordered in both the excited state (Fig. 2D) and the DNA-bound state(*33*), likely because of the helix-breaking Pro57 residue at position 2 of the putative hinge helix. In contrast, the rest of the protein from T11-Q56 undergoes significant changes in chemical shift upon transitioning to the excited state with an average CSP of 0.56 ± 0.27 ppm over 46 residues, underscoring the fact that disordered native CytR acquires a globally folded structure in the excited state. The magnitude of CSPs plotted on the structure of DNA-bound CytR^N^ (Fig. S5) also illustrate that CSPs are not localized to a particular region but instead distributed across the entire protein sequence from T11 to Q56.

Chemical shifts can also be used within the Random Coil Index framework to obtain estimates of generalized squared order parameter (S^2^) values (*43*) that are measures of the amplitude of fast dynamics occurring at a particular site in the protein. Figure 2E shows the residue-specific S^2^ values of the excited state evaluated from ^15^N, ^1^H^N, 13^Cα and ^13^CO chemical shift information. S^2^ values are high in the interior of CytR^N^ and range between 0.60 to 0.88, consistent with the rigidity expected from a globally folded excited state conformation. On the other hand, S^2^ values drop to less than 0.5 for the flexible terminal residues between M1-M12 and P57-E66 (Fig. 2E). Taken together, CSPs and S^2^ values demonstrate that natively disordered CytR^N^ undergoes a disorder-to-order transition, highlighting the novel ability of IDP sequences with a high net charge and a small fraction of hydrophobic residues to encode rigid globally structured topologies, and incurring a relatively small (ΔG_0,DE_ = 1.3 kcal/mol) free energy penalty.

### The excited state is a three-helix bundle containing a helix-turn-helix motif

In order to elucidate the atomic-resolution structure of the excited state, we set out to measure ^15^N-^1^H residual dipolar couplings (RDCs) (*44, 45*) as additional structural restraints for use in a structural calculation algorithm. ^15^N-^1^H RDCs were measured using either ^15^N-(*46*) or ^1^H^N^-CEST pulse sequences (Fig. S6). Briefly, in the ^15^N-CEST sequence, doublets are obtained by removing the 90_x_º-240_y_º-90_x_º ^1^H decoupling module during the exchange duration and the spacing between the two components of the doublet corresponds to the scalar coupling (^1^J_NH_) in the isotropic sample (Fig. 3A,top) and the sum of ^1^J_NH_ and the ^15^N-^1^H RDC in the aligned sample (Fig. 3A,bottom). Accordingly, the difference in doublet spacings between the two samples provides the value of the RDC (Fig. S6). On the other hand, the ^1^H^N^-CEST sequence is already designed so that every CEST profile represents a difference of the TROSY (H_z_N^β^) and anti-TROSY (H_z_N^α^) CEST profiles. Therefore, fitting each intensity dip to a difference of two Lorentzians directly furnishes ^1^J_NH_ for data acquired on an isotropic sample (Fig. 3B,top), or ^1^J_NH_+^15^N-^1^H RDC (Fig. 3B,bottom) in the case of an aligned sample (Fig. S6).

**Fig. 3.**
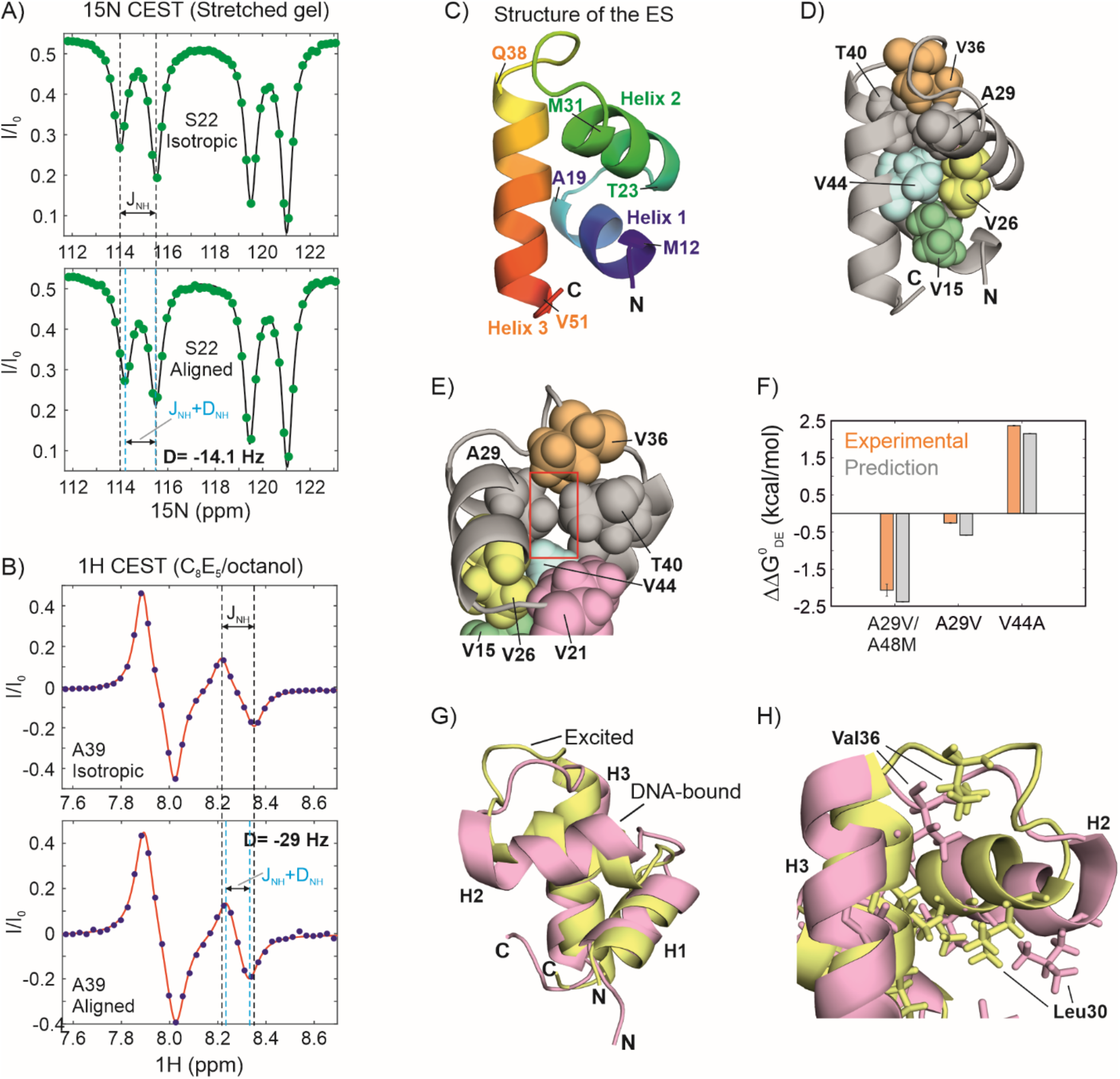
Structure of the CytR^N^ excited state. ^1^H-^15^N residual dipolar couplings (RDCs) for S22 (A) and A39 (B) of CytR^N^ obtained via a ^15^N CEST experiment lacking the ^1^H decoupling module during T_ex_ (A) or a ^1^H CEST experiment (B). CEST profiles were acquired on CytR^N^ aligned in stretched polyacrylamide gels (A) or in a lyotropic phase made from a C_8_E_5_/octanol mixture (B). In both panels CEST profiles at the top are from samples in isotropic media, while those at the bottom correspond to aligned samples. The black dashed lines indicate the position of the in-phase (A) or ‘anti-phase’ (B) doublets separated by ^1^J_NH_ in isotropic solution. Cyan lines reflect the positions of the same doublets in aligned media and are separated by ^1^J_NH_+^15^N-^1^H RDC. The RDC values in panels A and B are −14.1 Hz and −29 Hz respectively. C-E) Different orientations of the three-helix bundle structure of the globally folded excited state. The amino acids marking each helix boundary are indicated in panel C. The hydrophobic core of the ES of CytR^N^ is shown as spheres on the cartoon representation of the three-helix bundle in panels D and E. The core is made up of four conserved Val residues V15, V26, V36 and V44, and augmented by A29 and T40. The cavity formed by the smaller A29 (compared to the larger Val in other family members) is indicated by a red box in panel E. F) Predicted (grey) and experimentally observed (orange) free energy differences between the disordered and excited states of CytR^N^ mutants shown as differences from the wt free energy as a bar plot. Predictions were generated using the FoldX software suite(*53*), while experimental values are from global fits of CEST profiles for A29V and V44A CytR^N^ (Fig. S11,S13), and from volumes of HSQC spectral resonances for A29V/A48M CytR^N^ (Fig. S12). G,H) A comparison of the structures of excited (yellow) and DNA-bound (pink) CytR^N^ (PDB ID: 2LCV (*33*)). The sidechains of Leu and Val residues are shown as sticks and the sidechains of Leu30 and Val36, which are located near the DNA binding interface and change significantly in orientation between the excited and DNA-bound states, are indicated.

^15^N-labeled CytR^N^ was aligned either in a 6 % stretched polyacrylamide gel (PAG, Fig 3A) (*47*) or in a lyotropic phase made from bicelles containing a mixture of C_8_E_5_ and n-octanol (Fig. 3B) (*48*). Using a combination of the above ^15^N- and ^1^H^N^-CEST based methods, ^15^N-^1^H RDCs were obtained for 34 residues in PAG and ranged from −14 Hz to 6 Hz, while 31 RDCs varying between −29 Hz and 18 Hz were extracted from CytR^N^ aligned in C_8_E_5_/octanol (Fig. S7, Tables S4 and S5).

We then calculated the structure of the excited state of CytR^N^ using CS-Rosetta(*49, 50*) incorporating 119 backbone chemical shifts and 65 ^15^N-^1^H RDCs as structural restraints (Supporting Text). A well-defined funnel is observed in the energy vs RMSD plot, confirming that the structure calculations have converged (Fig. S8). Figure S8C shows the final ensemble of 10 excited state structures, defined by an all-atom RMSD of 0.84 Å and a Cα RMSD of 0.15 Å. There is good agreement between the input chemical shifts and the chemical shifts predicted by Sparta+ (*51*) for the excited state; the scatter in the correlations is of the same order of magnitude as the prediction accuracy of Sparta+ for each nucleus (Fig. S9). Moreover, RDCs predicted using the lowest energy structure agree very well with the experimental RDCs and return Q values of 0.124 and 0.091, and RMSD values of 0.87 Hz and 1.3 Hz for PAG and C_8_E_5_/octanol respectively (Fig. S10). In summary, the structure calculated using CS-Rosetta is a faithful representation of the NMR spectroscopic data collected on the CytR^N^ excited state.

CytR^N^ adopts a three-helix bundle topology in the excited state, in which helix 3 (H3) docks on to the helix-turn-helix (HTH) motif formed by helices 1 (H1) and 2 (H2) (Fig. 3C). The hydrophobic core is composed of five conserved valine residues V15, V21, V26, V36 and V44, supplemented by the methyl sidechains from A29 and T40 (Fig. 3D). Interestingly, while T40 is conserved within the LacR family, the residue at position 29 is occupied by a valine in FruR, PurR and LacR(*52*). The alteration of Val to Ala in CytR^N^ causes a small cavity in the excited state hydrophobic core (Fig. 3E) and CEST data (Fig. S11) show that the excited state of CytR^N^ is slightly stabilized by the A29V mutation, in which the larger Val sidechain can fill the hydrophobic core without generating cavities in the structure. When the core is further strengthened by replacing the small A48 sidechain with Met (the corresponding residue in LacR) (*52*), there is a dramatic stabilization of the excited state which becomes the dominant state in A29V/A48M CytR^N^ (p_E_ = 78 ± 6 %, Fig. S12), while the population of the disordered state is 22 ± 6 %. ^15^N and ^1^H^N^ chemical shifts of the ground state of A29V/A48M CytR^N^ match very well with the excited state chemical shifts of wt CytR^N^ (Fig. S12), establishing that this double mutant undergoes population switching. In contrast, disrupting the hydrophobic core by modifying V44 to the smaller Ala sidechain significantly destabilizes the excited state, which disappears from the CEST profile altogether (Fig. S13). The ΔΔG_0,DE_ values obtained from HSQC and CEST data for all three mutants agree well with structure-based stability predictions from FoldX (*53*) (Fig. 3F).

There is a close resemblance between the backbone secondary structures of the excited state and the DNA-bound form of CytR^N^ (*33*) (Cα backbone RMSD = 2.4 Å) (Fig. 3G), which is reflected in the excellent agreement between the Cα chemical shifts of the two states (Fig. S14A). However, there is more scatter in the correlation between the ^15^N (Fig. S14B) and ^1^H^N^ (Fig. S14C) chemical shifts of the excited and DNA-bound states; this is likely a consequence, not only of the charged DNA molecule containing aromatic nucleobases in the vicinity of DNA-bound CytR^N^, but also of small differences in helix orientation and hydrophobic packing between the two structures. For example, in DNA-bound CytR^N^ (*33*), the sidechain of Leu30 rotates outward towards the bound DNA, while Val36 moves inward towards the hydrophobic core, consistent with the tertiary structure of CytR^N^ adapting to the presence of the bound DNA molecule (Fig. 3H). In addition, the recognition helix H2 is slightly smaller in the excited state than in DNA-bound CytR and LacR.

### CytR^N^ binds DNA through the excited three-helix bundle state

The coexistence of the disordered ensemble of CytR^N^ with the three-helix bundle excited state, which is structurally similar to the DNA-bound state, raises the intriguing question of whether this excited state is selected by DNA molecules for binding. While this would result in a conformational selection (CS) mechanism of molecular recognition, the other possible models include the induced fit (IF) (*54*), which is known as the folding-upon-binding mechanism in IDP literature (*15*), and the triangular model, in which binding occurs through both the CS and IF pathways.

We addressed this question by first choosing D34 as a simultaneous NMR reporter of the disordered state (D), the excited conformation (E) and the DNA-bound form (B). D34 is located in the H2-H3 loop and has distinct chemical shifts of 119.6 ppm, 113.9 ppm and 117.2 ppm in D, E and B respectively. ^15^N CEST profiles of the D34 resonance in the absence of DNA show two intensity dips at the chemical shifts of the disordered (D) and excited (E) states (Fig. S15). We then acquired CEST data on a sample containing 628 µM ^15^N-labeled CytR^N^ and 150 µM uridine phosphorylase (*udp*) half-site double-stranded DNA, which is one of the operator regions specifically recognized by CytR^N^ (*33*). In the presence of cognate DNA, a distinct third dip in intensity can be clearly discerned in the CEST profile of CytR^N^ D34 at the ^15^N chemical shift of state B (117.2 ppm) which is the direct result of exchange between free and DNA-bound forms of CytR^N^ (Fig. 4A).

**Fig. 4.**
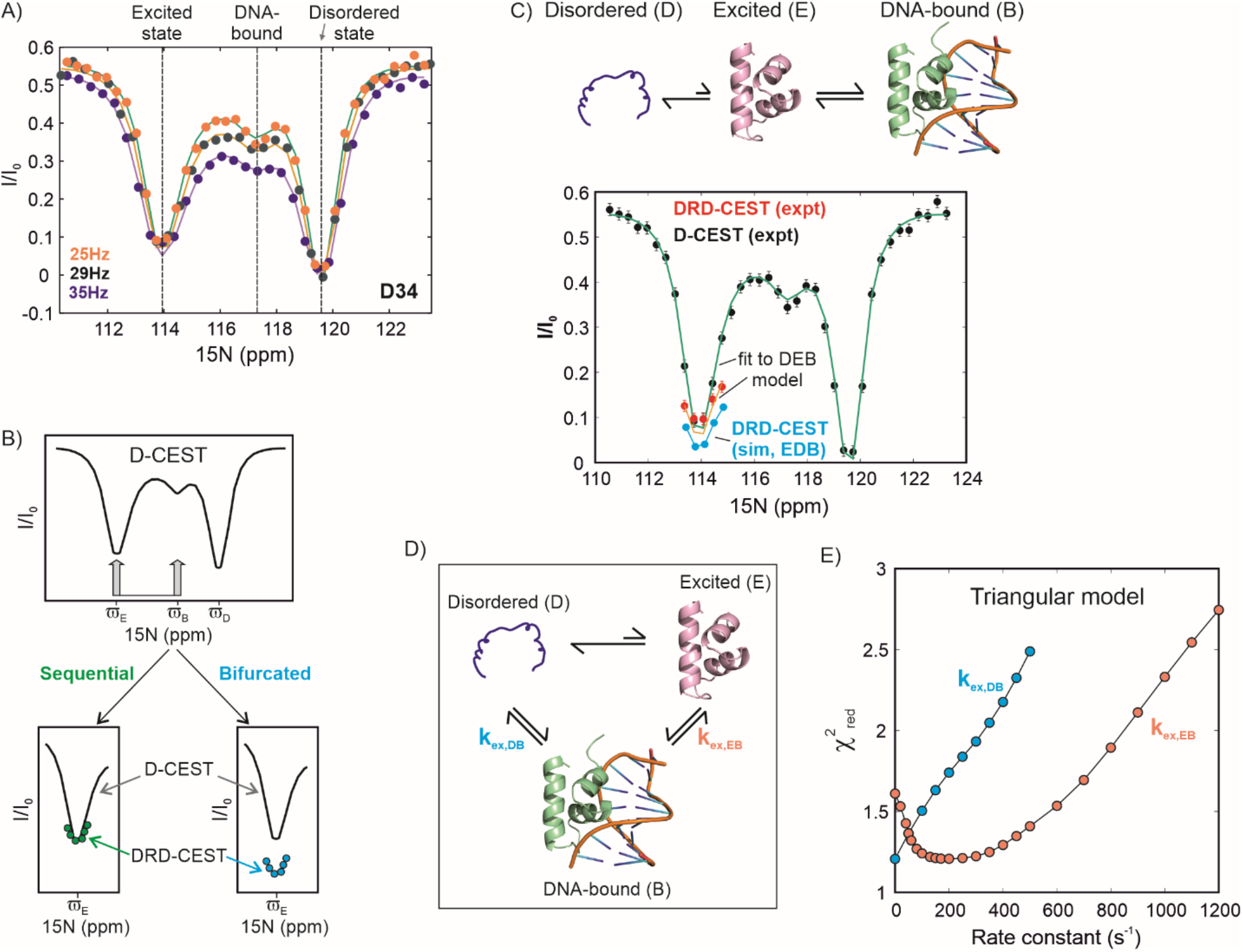
The excited state of CytR^N^ is responsible for functional regulation. A) ^15^N CEST profiles of D34 in the presence of DNA acquired with B_1_ field strengths of 25, 29 and 35 Hz on a sample containing 628 µM CytR^N^ and 150 µM DNA. The three dips in intensity at the chemical shifts of the disordered (119.6 ppm), excited (113.9 ppm) and DNA-bound (117.2 ppm) states are indicated with dashed lines. B) Schematic representation of the double resonance DANTE-CEST (DRD-CEST) experiment (*55*) for a system undergoing three-state exchange between one visible (D) and two invisible states (E and B). The D-CEST profile for this system is shown at the top. Overlays of the minor dip at µ_E_ in D-CEST (black line) and DRD-CEST (coloured circles) are shown at the bottom for a sequential (left) and a bifurcated model (right). C) ^15^N CEST (black circles) and DRD-CEST (red circles) profiles of D34 acquired on the same sample as panel A. Green and orange lines are global fits of the regular and DRD-CEST data at multiple B_1_ fields to the D↔E↔B model shown above the CEST profile. Cyan circles are simulated data DRD-CEST data points assuming a bifurcated E↔D↔B model. Simulations were done using best-fit parameters obtained by fitting 5 D-CEST and 3 DRD-CEST datasets globally to the E↔D↔B model. D) The triangular model for binding of CytR^N^ to DNA, where rate constants describing the binding of D and E are k_ex,DB_ and k_ex,EB_ respectively. (E) 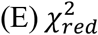 distribution of k_ex,DB_ (cyan) and k_ex,EB_ (orange) generated from fits of regular and DRD-CEST data to the triangular model.

Having obtained CEST profiles for D34 which report on the simultaneous exchange between D, E and B, we turned to the recently developed Double Resonance DANTE-CEST (DRD-CEST) (*55*) for determining whether binding occurs via the excited state. DRD-CEST has demonstrated potential for unequivocally distinguishing between sequential and bifurcated models of chemical exchange(*55*). In the case of CytR^N^, viewed from the perspective of the NMR-visible disordered state D, the CS mechanism of binding (D↔E↔B) is a sequential model while the IF pathway is a bifurcated model (E↔D↔B). DRD CEST builds upon D-CEST or DANTE CEST(*56*), in which the continuous wave irradiation that is typically employed during the CEST exchange time is replaced by the DANTE selective excitation scheme (*57*) (Fig. 4B). When implementing DRD-CEST, a sufficiently large B_1_ field is first chosen so that the magnetization of molecules arriving at E are completely dephased with respect to the starting D magnetization (Fig. 4A). Then, the sample of CytR^N^ containing DNA is simultaneously irradiated with this B_1_ field at the chemical shifts of both E and B, and the CEST profile of D34 is quantified. If the model of conformational exchange is a sequential one, irradiation at E and B will not change the size of the dip compared to irradiation only at E. This is because dephasing is complete at E itself, so additional irradiation at D does not make a difference. On the other hand, if the model is bifurcated, dephasing at E and dephasing at B occur on magnetization from different molecules, so even if dephasing is complete at E, simultaneous irradiation at E and B will increase the dip size compared to irradiation only at E. Therefore, if conformational exchange proceeds through a sequential model, the size of the dip at the chemical shift of state E remains unchanged between the regular and DRD-CEST profiles. On the other hand, the fingerprint of a bifurcated model is a pronounced increase in dip size in the DRD-CEST profile compared to regular CEST data (Fig. 4B).

Figures 4C and S16 show the regular CEST profile of D34 (D state) in black and the DRD-CEST data in red. The size of the intensity dip for state E clearly remains the same between the CEST and DRD-CEST profiles, unambiguously establishing that DNA binding occurs via the excited state. The simulated dip for a bifurcated model is shown in cyan circles for comparison. In order to further evaluate the binding mechanism, we globally fit the CEST (5 B_1_ fields) and DRD-CEST data (3 B_1_ fields) separately to the sequential and bifurcated models. The 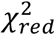 value for the bifurcated model 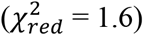 is substantially higher than the sequential model 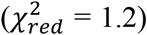, signifying that the CEST data are better described by the CS mechanism (Fig. S16).

Finally, we quantitatively assessed whether both the CS and IF mechanisms are simultaneously operative in the DNA-CytR^N^ interaction by fitting the CEST and DRD-CEST data globally to a triangular model (Fig. 4D); here, k_ex,DB_ (=k_DB_+k_BD_) and k_ex,EB_ (=k_EB_+k_BE_) are the rate constants for the binding of states D and E to DNA respectively. The best-fit values of k_ex,DB_ and k_ex,EB_ from the triangular fit are 0.00015 ± 0.3 and 194 ± 51 s^-1^ respectively, signifying that the unfolded state cannot directly bind DNA to form the specific DNA-CytR^N^ complex. 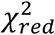 surfaces (Fig. 4E) show that the optimal fit of the triangular model to the CEST data is obtained when k_ex,DB_ is very close to 0. In contrast, there is a heavy penalty in 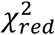 when the k_ex,EB_ is forcibly set to 0 and that the optimum value is around 200 s^-1^, which matches the k_ex,EB_ value of 194 s^-1^ obtained above. A flux-based analysis (*58*) of the triangular model (Supporting Text) shows that the flux along the CS pathway is at-least ∼200-fold (and up to ∼10^6^-fold) larger than the flux along the IF pathway.

In summary, the CEST and DRD-CEST data unequivocally establish that the DNA recognition pathway of disordered CytR^N^ predominantly follows a folding-before-binding mechanism, where molecules of CytR^N^ in the structured excited state but not the disordered state bind to DNA (Fig. 5).

**Fig. 5.**
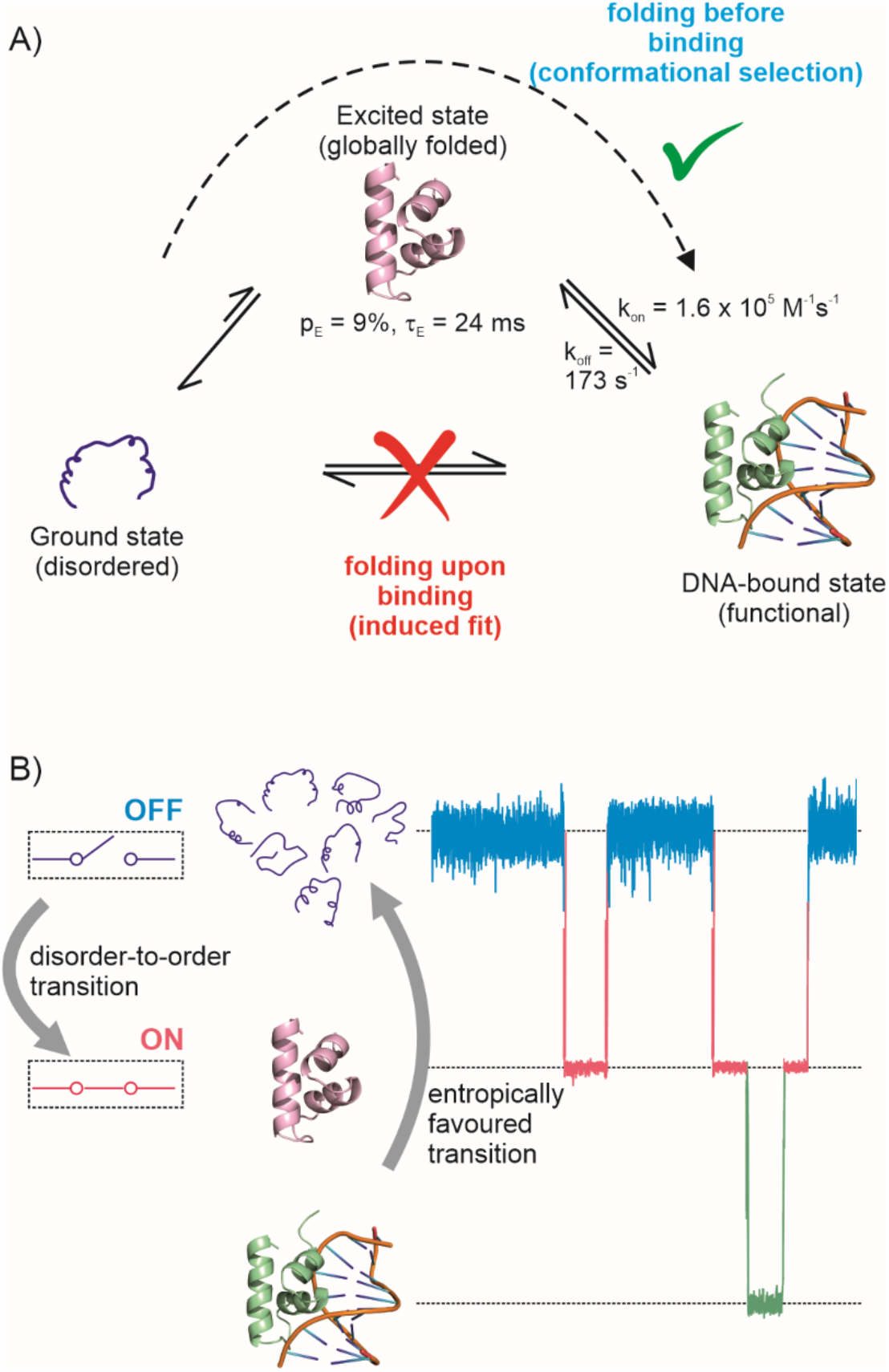
The ‘folding-before-binding’ model for DNA recognition by intrinsically disordered CytR^N^ as a regulatory switch. A) CytR^N^ is intrinsically disordered in its native state and the molecular recognition event leading to the DNA-bound functional form involves a conformational selection-based folding-before-binding mechanism. B) A model for how the disorder-to-order transition in CytR^N^ turns on the regulatory switch into the functional state and primes it for DNA binding, while the large entropy of the disordered state favours the dissociation of the complex and subsequent unfolding of the folded conformation, resetting the switch into the ‘off’ mode. Molecular dynamics traces of CytR^N^ are schematic representations and do not correspond to actual data.

## Discussion

How intrinsically disordered proteins function despite lacking stable secondary and tertiary structure remains an open question. In this work, we have demonstrated that intrinsically disordered CytR^N^, which belongs to the iconic LacR family of bacterial transcriptional repressors, transiently adopts a globally folded structure that is accessible via thermal fluctuations. We have also provided conclusive evidence that this disorder-to-order transition functions as a regulatory switch by converting the binding-incompetent disordered ensemble into a three-helix bundle HTH structure capable of binding DNA. In stark contrast to the IF folding-upon-binding models that are routinely used to describe molecular recognition by IDPs, the CytR^N^-DNA interaction follows a CS mechanism that we term ‘folding-before-binding’ (Fig. 5).

The free energy landscape of an IDP controls its folding and molecular recognition properties and is often depicted as a flat surface with multiple shallow minima separated by very small barriers(*4–6*). Interconversion between these structurally similar conformations occurs on the picosecond-nanosecond timescales. Kinetically distinct higher free energy excited states in the IDP landscapes are considered to be thermally inaccessible because they are significantly destabilized with respect to the high entropy disordered native ensemble(*4*). While this may generally be true, intrinsically disordered CytR^N^ furnishes a striking counter-example by populating a globally ordered three-helix bundle that is a mere 1.3 kcal/mol (2.4 kT) higher in free energy than the disordered ensemble. This excited state is not stabilized by interaction with partner ligands, osmolytes, crowding agents or post-translational modifications, but is instead as much an intrinsic property of the free energy landscape of CytR^N^ as the native disordered ensemble. Moreover, the detection of an excited state of CytR^N^ with a millisecond lifetime confirms that IDPs can dynamically sample conformations over a hierarchy of timescales ranging from picoseconds to milliseconds. In the case of CytR^N^, this slow disorder-to-order transition has been leveraged for performing its DNA-binding function.

The coexistence of the disordered and three-helix bundle conformations of CytR^N^ is eventually encoded in its amino acid sequence. IDP sequences are characterized by low hydrophobicity, a high net charge and a paucity of large aromatic and aliphatic hydrophobic amino acids(*1*), and the sequence of CytR^N^ meets all these criteria. Our results therefore clearly show that sequences classified as IDPs can nevertheless accommodate not only locally folded, but also globally ordered regions at comparable free energies to the disordered state. In our case, the precise sequence determinants which allow the *cytR* gene to encode a thermally accessible excited state remain tantalizingly unknown. However, a comparison of the sequences of CytR^N^ with its family members FruR, LacR and PurR (*52*) shows that CytR retains most of the hydrophobic core necessary for forming the three-helix bundle, while increasing the fraction of positively charged residues in the variable regions of the sequence. Simultaneously, two large hydrophobic residues at sites 29 and 48 are replaced with Ala in CytR^N^, resulting in a slight loosening of the hydrophobic core and concomitant destabilization(*52*). The synergistic combination of the above factors seems to assist CytR^N^ in retaining its intrinsically disordered state without rendering the folded state thermally inaccessible. Indeed, our data clearly shows that the key A29V/A48M mutant swaps the folds of the ground and excited states in CytR^N^, making the folded three-helix bundle conformation more stable than the intrinsically disordered enesmble.

Our CEST data shows the formation of the specific CytR^N^-DNA complex occurs primarily by the binding of DNA to the globally structured excited state. Since CytR^N^ is disordered in its native state and folded in the bound form, the CytR^N^-DNA interaction can be still classified as folding coupled to binding. However, unlike conventional IF-like folding-upon-binding situations that are prevalent in IDP literature (*14, 15, 17*), CytR^N^ folds before binding DNA and therefore conforms to a classic CS mechanism. There are a number of points that are noteworthy in this regard. First, the function of the intrinsically disordered CytR^N^ domain is to bind DNA. Since this function is carried out by the three-helix bundle folded conformation, the excited state is responsible for the functional regulation of the CytR^N^ IDP. The overall phenomenon of transcriptional repression at the promoter and operator sites involves the C-terminal domain of CytR^N^, other proteins like CAP (catabolite activator protein) and the inducer cytidine, all of which modulate the binding of DNA to CytR^N^ (*25*). Nevertheless, our conclusions are concerned with the elementary functional binding reaction of CytR^N^ with DNA and affirm that only the excited state and not the disordered form is capable of forming the specific DNA-CytR^N^ complex. Second, our classification of the mechanism as CS pertains only to global overall changes in structure between the free and DNA-bound forms of CytR^N^; they do not rule out small IF-like adjustments of sidechains in response to DNA binding. Indeed, the orientation of sidechains proximal to the DNA binding site, such as L30 and V36 are different between DNA-bound and excited CytR^N^, suggesting that the HTH motif subtly reorganizes itself in the complex to accommodate the DNA molecule (Fig. 3H). Interestingly, additional evidence for IF-like behaviour is seen near the end of the recognition helix H2, where the terminal N25 forms hydrogen bonds with the bound DNA in the structure of the LacR/operator DNA complex (*59, 60*), but the corresponding residue N32 is oriented differently in the CytR^N^ excited state (Fig. S17A). Unlike for most of CytR^N^, CEST data (Fig. S17B) demonstrate that the chemical shifts of N32 are significantly different in the DNA-bound and excited conformations, reflecting the IF-like alterations in CytR^N^ structure occurring after the DNA binding event. Third, our observation of a CS mechanism in the case of molecular recognition by CytR^N^ raises the intriguing question of what determines whether an IDP will adopt a folding-before-binding or a folding-upon-binding pathway. We hypothesize that a folding-before-binding route is likely for IDPs where the bound form exhibits a globally folded conformation with substantial secondary structure and tertiary contacts. On the other hand, IDPs which form short helices connected by linear unstructured polypeptide segments in the bound state may require more assistance from the partner protein for stabilizing the local secondary structural elements and therefore adopt a folding-upon-binding pathway. We also speculate that a number of instances of folding-before-binding may have been missed in literature, simply because the excited folded conformations are transiently and sparsely populated, and therefore invisible to most biophysical methods. Finally, the CEST data clearly indicate that there is no measurable kinetic coupling between the disordered ensemble and the specific DNA-CytR^N^ complex, and that virtually all the flux to this bound state traverses through the excited conformation. This rules out models that involve a rapid conformational search by the disordered state and subsequent folding on the DNA once the specific operator sequence is located. However, models where the unstructured state scans the DNA by 1D diffusion, dissociates transiently to adopt the folded conformation and then binds to the specific operator site cannot be excluded by our data, since binding in such models eventually occurs via the excited state.

The existence of a novel mode of functional regulation of a bacterial IDP by a conformationally excited state is pertinent because bacteria, unlike eukaryotes, are unable to expand the structural and functional repertoire of their proteomes through post-translational modifications. Bacteria also appear to be limited in their ability to use membraneless condensates for spatiotemporal control of cellular processes(*61*). It is, therefore, appropriate that a bacterial IDP relies on a more primitive disorder-to-order regulatory switch that operates simply by fine-tuning the balance between hydrophobicity and intrinsic net charge to eventually modulate the conformational free energy surface; in-turn, this suggests that dynamic regulatory pathways may be more prevalent in bacterial IDPs as a way to expand the functionality of their limited genetic material. Additionally, HTH motifs are ubiquitous nucleic acid recognition modules that have diversified enormously in the architectural context in which they are found, as well in the function they perform in the cell(*62*). The appearance of a folding-before-binding mode of DNA recognition in a HTH-containing bacterial protein suggests that this mechanism was available early on in evolution and might have become incorporated into intrinsically disordered transcriptional repressors in higher organisms as well.

Although the exact role of disorder in the DNA binding domain of CytR is yet to be deciphered, we adapt a model proposed in literature (*63, 64*), in which the disordered state serves to reset the CytR^N^ regulatory switch (Fig. 5B). Since natively disordered CytR^N^ has a large chain entropy, folding and subsequent complex formation with DNA will incur a free energy cost, which has been estimated in literature to be of the order of 2.5 kcal/mol (*65*). Studies on the c-Myb/CBP-KIX system show that the extent of disorder is correlated with k_off_ values (*66*), implying that this 2.5 kcal/mol free energy penalty associated with disorder in CytR^N^ could decrease the lifetime of the CytR^N^-DNA complex and offer an attractive route to turn off the regulatory switch. Thus, while the disorder-to-order transition ‘turns on’ the regulatory switch and primes it to bind DNA, the disorder in the native state drives the switch towards the ‘off’ position.

Finally, the dynamic regulatory switch in CytR^N^ is another example of molecular recognition occurring via a thermally accessible excited state. It emphasizes the point that dynamics is pivotal for function and that a comprehensive understanding of the relationship between structure and function cannot be obtained merely by studying the native state alone. In the context of IDPs, a thorough characterization of the free energy surface is particularly vital, not only because IDPs lack the stable structure traditionally deemed to be necessary for function, but also because they have recently gained importance as targets for pharmacological intervention (*67*).

## Acknowledgments

The authors thank Lewis E. Kay (University of Toronto) for providing the pulse sequence codes used in this work, Nikolaos Sgourakis, Santrupti Nerli, Jayashree Nagesh and Sandhya Sankaran for help with CS-Rosetta, and Lewis Kay, Pramodh Vallurupalli, Siddhartha Sarma, Bharathwaj Sathyamoorthy and Rina Rosenzweig for stimulating discussions.

## Funding

DBT/Wellcome Trust India Alliance Fellowship IA/I/18/1/503614 (A.S.)

DST/SERB Core Research Grant CRG/2019/003457 (A.S.)

Start-up grant from the Indian Institute of Science Bangalore (A.S.)

Infrastructural support from the following programs of the Government of India: DST-

FIST, UGC-CAS, and the DBT-IISc partnership program (A.S.)

Fellowship support from CSIR (K.M.)

Fellowship support from the Indian Institute of Science Bangalore (B.N.)

## Author contributions

Conceptualization: ANN, AS

Methodology: KM, AS

Investigation: KM, BN, SM, ANN, AS

Visualization: KM, BN, AS

Funding acquisition: ANN, AS

Project administration: ANN, AS

Supervision: AS

Writing – original draft: AS

Writing – review & editing: KM, BN, SM, ANN, AS

## Data and materials availability

The backbone resonance assignments of wt CytR^N^ are deposited in the Biological Magnetic Resonance Databank (BMRB) (accession number: 51449). For raw data and materials requests, please contact A.S.

## Supplementary Materials

Materials and Methods

Supplementary

Text Figs. S1 to S17

Tables S1 to S6

References (68 to 78)

