## Supporting Information for "Functional regulation of an intrinsically disordered protein via a conformationally excited state"

### Materials and Methods

#### Overexpression and purification of isotope-labeled proteins for NMR spectroscopy

U-$^{15}N$ labeled CytR. Full-length CytR was cloned between the NdeI and HindIII sites in the pET-29b(+) vector along with an N-terminal TEV-cleavable 6x His-tag, followed by a linker. *Escherichia coli* (*E. coli*) BL21(DE3) cells were used for protein expression and $^{15}N$ isotope labelling was carried out by growing cells in M9 minimal media containing 1g/L $^{15}NH_4Cl$. Cells were grown at 37 ºC till an OD of 0.8, induced with 1 mM isopropyl β-D-1-thiogalactopyranoside (IPTG) and then shaken at 20 °C for ~ 20 hours. Cells were harvested by centrifugation (at 4 °C and 13,000 rpm) and the cell pellet were resuspended in Buffer A (20 mM TRIS (pH=7.5), 1 mM BME, 1 mM EDTA and 300 mM NaCl) containing 2 mM $MgCl_2$, and a Roche protease inhibitor tablet. Cells were lysed with lysozyme (7.5 mg/g of cell pellet) and sonication, following which the suspension was incubated with PMSF (1 mM final concentration), DNase I (50 mg/g of cell pellet) and polyethyleneimine (0.04 % of the total volume) at room temperature for one hour. The suspension was then centrifuged for 30 min at 13,000 rpm and 4°C, and the clarified lysate was loaded on a Ni-NTA column. His-tagged protein was eluted from the column using Buffer A containing 250 mM imidazole and the tag was subsequently cleaved using TEV protease while simultaneously dialyzing against Buffer A at room temperature. Tagless CytR was then recovered by passing the solution again through the Ni-NTA column. Flow-through and wash fractions containing CytR were concentrated and purified using size-exclusion chromatography on a Superdex 200 column equilibrated in 150 mM ammonium acetate. Pure fractions were checked with SDS-PAGE on a 15 % gel, pooled, dialyzed into 22 mM sodium phosphate buffer (pH 7), flash frozen and stored at -80 ºC.

U-$^{15}N$ and U-$^{15}N$,$^{13}C$ labeled CytR$^N$. Untagged CytR$^N$ (CytR$_{1-66}$) was cloned between the NdeI and HindIII sites in the pET-29b(+) vector. Overexpression was carried out as described above for full-length CytR. U-$^{15}N$ and U-$^{15}N$,$^{13}C$ isotope-labeled CytR$^N$ were obtained from cells grown in M9 minimal media supplemented with 1 g/L $^{15}NH_4Cl$ without (U-$^{15}N$) or with 3 g/L $^{13}C_6$-glucose (U-$^{15}N$,$^{13}C$). CytR$^N$ was purified using a previously described protocol with small modifications(*33*). Briefly, the clarified cell lysate was subjected to 60 % ammonium sulphate precipitation. The precipitate was removed by centrifugation and the supernatant containing CytR$^N$ was loaded on an SP-sepharose cation exchange column. CytR$^N$ elutes at an ionic strength of 540-640 mM. Fractions containing CytR$^N$ were pooled, concentrated and purified on a Superdex 75 size exclusion column, from which CytR$^N$ elutes at a volume of ~ 70 mL. Fractions were checked for purity using SDS-PAGE, and pure CytR$^N$ was dialyzed against 150 mM ammonium acetate, lyophilized and stored at -20 ºC.

U-$^{15}$N,$^{13}$C A29V/A48M, U-$^{15}$N A29V and U-$^{15}$N V44A CytRN. A29V and V44A mutations were generated in the above construct of CytR$^N$ by site-directed mutagenesis using the method of overlapping primers. For A29V/A48M CytR$^N$, the A29V single point mutation was introduced using site directed mutagenesis in CytR$^N$ cloned in the pTXB1 vector and the A48M mutation was subsequently introduced in the A29V CytR$^N$ mutant. Isotope-labeled mutant CytR$^N$ variants were overexpressed and purified using the same protocol as for wild-type CytR$^N$. U-$^{15}$N,$^{13}$C A29V/A48M CytR$^N$ was overexpressed in minimal media as described above and the purification was carried out as reported previously (*52*).

Oligonucleotide preparation. Single-stranded *udp* half-site DNA (ssDNA) sequences were purchased from Sigma (5′-ATTTATGCAACGCA-3′). Forward and reverse ssDNA were dissolved in 22 mM sodium phosphate buffer pH 7 and their concentrations were estimated using UV absorbance at 260 nm. A 1.1-x buffer was used for dissolving the DNA in order to account for the 10 % $D_2O$ added as NMR lock solvent. Equimolar concentrations of forward and reverse DNA strands were mixed and the DNA was annealed by first incubating at 95 °C for 15 minutes on a dry bath, and then lowering the temperature slowly by turning off the dry bath. The double stranded DNA was stored at -20 °C for further use.

**Sample preparation**

All NMR experiments were carried out on samples in 20 mM sodium phosphate buffer (pH 7.0) containing 10 % $D_2O$ for the field-frequency lock. Sample concentrations ranged from 60 μM (full-length CytR) to 2 mM. The details of the various CytR$^N$ samples used are summarized in Table S1.

Sample preparation for RDC measurements

Polyacrylamide stretched gels**:** 6% polyacrylamide gels were cast using a 30 % acrylamide: bisacrylamide (29:1) mixture made in Tris buffer pH 8.8 and polymerized in a 6 mm gel casting chamber (www.newera-spectro.com). After the gel solidified (~ 15 minutes), the gel was dialysed multiple times against water to remove the buffer components and air-dried. Before making RDC measurements, the gel was soaked in 550 µL of protein solution containing 10 % $D_2O$ for about 10-12 hours using the same casting chamber. This gel was then transferred to a 4.2 mm (inner diameter) RDC NMR tube with the help of a piston driver. 20 µL of buffer was added to both ends to prevent drying of the gel and the bottom of the sample tube was sealed with a gel end plug. At the top, the gel was capped with a support rod carrying a top-gel plug at the sample end. The support rod was held in place with a support cap that attaches to the NMR tube.

$C_8E_5$/n-octanol: Prior to preparing the protein sample, a 10 % $C_8E_5$/n-octanol mixture was prepared in the following manner (*48*): 36 µL of $C_8E_5$ (Sigma) was dissolved in 222 µL of sodium phosphate buffer (pH 7) and 30 µL $D_2O$. The $C_8E_5$ solution was kept on ice throughout and mixed thoroughly. 12 µL n-octanol was added to the above $C_8E_5$ solution in aliquots of 4 µL and the mixture was vortexed after each addition. The solution went from yellow to turbid and finally a clear viscous lyotropic phase was obtained. A $^2H$ 1D spectrum was recorded on this sample and a residual quadrupolar splitting from the $^2H$ nucleus of HDO molecules of 43 Hz was observed. Subsequently, 300 µL of the same buffer lacking the protein was added to dilute the mixture to 5 % $C_8E_5$.

A similar 10 % $C_8E_5$/octanol mixture was then prepared and diluted to 5% with buffer containing 1.6 mM $^{15}N$-labeled $CytR^N$ (Sample 7). RDC measurements were collected on this sample as described below. The $^2H$ residual quadrupolar splitting for the final sample was 30.2 Hz.

**NMR data collection and analysis**

NMR data collection and processing. All NMR experiments were carried out at 15 ºC using a 14.1 T ($^1H$ Larmor frequency of 600 MHz) Agilent DD2 spectrometer equipped with cryogenically cooled or room-temperature single-axis gradient triple resonance probes or a 16.4 T ($^1H$ Larmor frequency of 700 MHz) Bruker Avance Neo spectrometer equipped with a room-temperature TXI single-axis gradient triple resonance probe. NMR spectra were processed using NMRPipe(*68*) and visualized using the NMRDraw(*68*) and NMRFAM-Sparky software packages(*69*).

Backbone resonance assignments of CytR$^N$. The backbone chemical shifts of wt (Sample 1) and A29V/A48M CytR$^N$ (Sample 12) were assigned using a 2D $^1$H-$^{15}$N HSQC and standard 3D triple resonance datasets HNCACB, CBCA(CO)NH, HN(CA)CO and HNCO (*70*). Assignments of wt CytR$^N$ were directly transferable to the $^1$H-$^{15}$N HSQC spectra of A29V (Fig. S11) and V44A CytR$^N$ (Fig. S13), as the chemical shift perturbations to the disordered state are very small (0.078 and 0.247 ppm in $^{15}$N and 0.004 and 0.07 ppm in $^1$H averaged over 55 residues for A29V and 51 residues for V44A CytR$^N$ respectively).

Chemical exchange saturation transfer (CEST) data acquisition. $^{15}$N, $^1$H$^N$, $^{13}$C$\alpha$, $^{13}$C' CEST data were acquired on U-$^{15}$N,$^{13}$C samples of CytR$^N$ (Samples 1 and 2) using pulse sequences reported in literature(*37-40*). The details of the experiments are summarized in Table S2. The $B_1$ field was calibrated using the method reported by Guenneugues *et al.* (*71*). CEST profiles were obtained as a plot of the offset frequency vs the ratio of the peak intensity ($I/I_0$) where I and $I_0$ are the intensities obtained in the presence and absence of an exchange duration in the CEST pulse sequence. $^{15}$N CEST profiles of A29V and V44A CytR$^N$ were acquired using Samples 10 and 11 respectively.

Excited state RDC measurements were made using separate isotropic and aligned samples for each alignment medium (Samples 4-7). $^{15}$N CEST data for RDC measurements were acquired by setting the power level of the $^1$H decoupling module during $T_{ex}$ to 0 W. $^1$H$^N$ CEST profiles were collected without any modification to the previously reported pulse sequence (*38*). Details of data acquisition are reported in Table S4.

DANTE-CEST (D-CEST) and Double resonance DANTE-CEST (DRD-CEST) data were acquired on DNA-bound CytR$^N$ (Sample 4) on a Bruker 700 MHz spectrometer. The DANTE train consisted of pulses applied at an RF field of 3.57 kHz. Double resonance DANTE-CEST (DRD-CEST) measurements were carried out by keeping DANTE sweep window ($sw_{DANTE}$) fixed to a value of 254 Hz, which corresponds to the separation in $^{15}$N frequency between the excited state and the DNA-bound state resonances of D34.

**Analysis of CEST profiles**

$^{15}$N CEST of wt CytR$^N$ without DNA. $^{15}$N CEST data acquired at 2 $B_1$ fields for wt CytR$^N$ were fit to the two-state Bloch-McConnell equations (*72*) using the software package ChemEx (https://github.com/gbouvignies/ChemEx). 12 residues, A16, A19, S22, A24, T25, N32, T40, R49, G52, Y53, L54 and Q56 were selected since they had isolated resonances in the $^{1}H$-$^{15}N$ HSQC that showed well-separated major and minor dips in CEST profiles and these residues were used to get estimates of $p_E$ and $k_{ex}$. Monte Carlo error distributions of $p_E$ and $k_{ex}$ were obtained by fitting 1000 replica datasets to the two-state Bloch-McConnell equations using the same fitting protocol described above. These replicas were constructed from the original CEST dataset by adding random noise with zero mean and the same standard deviation as the error in $I/I_0$, assuming that the noise originates from an underlying Gaussian distribution. $\chi^2_{red}$ plots for $p_E$ ($k_{ex}$) were constructed by keeping the value of $p_E$ ($k_{ex}$) fixed at various during the fitting routine and repeating the fit.

For some of the overlapped resonances, the overlap was resolved at 25 ºC and CEST profiles acquired at 25 ºC were used to assign excited state chemical shifts to each of the overlapped residues. In a subset of these cases, CEST profiles showed two minor dips, one originating from each of the two overlapped peaks. In such cases, CEST profiles were fit to a sum of Lorentzians in order to extract residue-specific chemical shifts of the excited state. Six residues were eliminated from the analysis because they were severely overlapped in the HSQC spectra and the overlap could not be resolved at higher temperature.

$^{15}N$ CEST A29V and V44A CytR$^{N}$. $^{15}N$ CEST profiles of A29V CytR$^{N}$ were fit to a two-state model of conformational exchange using ChemEx as described for wt CytR$^{N}$. 12 residues were chosen for the global fitting routine in order to determine $p_E$ and $k_{ex}$. Since only one intensity dip was present in all CEST profiles of V44A CytR$^{N}$, these CEST profiles could not be fit to the two-state Bloch-McConnell equations. Instead, an upper limit to the population of the excited state was estimated as described in the legend to Figure S13.

$^{13}C\alpha$, $^{13}C'$ and $^{1}H^{N}$ data. $^{13}C\alpha$, $^{13}C'$ and $^{1}H^{N}$ chemical shifts of the CytR$^{N}$ excited state were extracted from the corresponding CEST profiles, either by fitting the data to a two-state exchange model within ChemEx, or by fitting the data to a sum of Lorentzians. In case of $^{1}H^{N}$ CEST, each dip is a difference of two Lorentzians separated by $^{1}J_{NH}$, which was kept fixed at a value of 93 Hz during the fitting procedure.

<u>$^{15}N$ and $^{1}H^{N}$ CEST data for measuring RDCs</u>. $^{15}N$ and $^{1}H^{N}$ CEST profiles acquired on isotropic and aligned samples were modeled as sums of Lorentzians, where each dip in $^{15}N$ ($^{1}H^{N}$) CEST was a sum (difference) of two Lorentzians. The frequency difference between the two Lorentzians for a dip gives the $^{1}J_{NH}$ value for the isotropic sample, and the $^{1}J_{NH}$+RDC value for the aligned sample.

<u>D- and DRD-CEST data</u>. D- and DRD-CEST data acquired on DNA-bound $^{15}N$ CytR$^{N}$ was modeled using three-state Bloch-McConnell equations within the ChemEx software package. D-CEST data collected at 5 $B_1$ fields (8.8, 15.9, 24.8, 28.7 and 34.7 Hz) and DRD-CEST data collected at 3 $B_1$ fields (25.0, 28.9, 34.9 Hz) for D34 were globally fit to either linear (D↔E↔B and E↔D↔B) or triangular models of chemical exchange. In each modeling protocol, all parameters except $k_{ex,DE}$ were allowed to float while $k_{ex,DE}$ was fixed to the value obtained in the absence of DNA. The excited state population ($p_E$) was not fixed to the value in the absence of DNA because there are indications in literature that the folded conformation of CytR$^{N}$ is stabilized by the electrostatic field of the DNA molecule. $\chi^2_{red}$ surfaces for $k_{ex,EB}$ and $k_{ex,DB}$ were constructed by fixing the respective parameters to specific values and determining the $\chi^2_{red}$ from the fit.

<u>Chemical shift perturbations (CSPs)</u>. CSPs between the excited state and the disordered conformation were calculated from CEST-derived $^{15}N$, $^{1}H^{N}$, $^{13}C\alpha$ and $^{13}C'$ chemical shifts for the excited state and assigned backbone shifts of the native state using the formula:

$$CSP_i = \sqrt{\frac{1}{N_i}\sum_{j}^{N_i}\left(\frac{\Delta\omega_{ij}}{\Delta\omega_{j,RMS}}\right)^2}$$

Here, $i$ is the residue index and $CSP_i$ are the residue-specific chemical shift perturbations. $N_i$ is the number of nuclei for the $i^{th}$ residue for which excited state chemical shifts are available and $j$ is the index that runs from 1 to $N_i$. $\Delta\omega_{ij}$ are the chemical shift differences between the excited and ground states for the $i^{th}$ residue and the $j^{th}$ nucleus while $\Delta\omega_{j,RMS}$ is 1 standard deviation of the spread in protein chemical shifts for the $^{15}N$ (6.10 ppm), $^{1}H^{N}$ (0.93 ppm), $^{13}C\alpha$ (2.94 ppm) or $^{13}C'$ (4.07 ppm) nuclei based on data deposited in the Biological Magnetic Resonance Databank (BMRB).

Secondary structural propensity (SSP). SSP calculations were performed using the SSP software package. While $^{13}C\alpha$, $^{13}C\beta$, $^{1}H^{N}$, $^{15}N$ and $^{1}H\alpha$ chemical shifts obtained from BMRB (accession number 17419) were used to determine SSP scores for the DNA-bound state, only $^{13}C\alpha$, $^{13}C\beta$, and $^{1}H\alpha$ chemical shifts were used for the disordered form (*32*).

### Supplementary Text

#### Structure calculations using CS-Rosetta

The structure of the excited state of $CytR^{N}$ was calculated using a standard CS-Rosetta structure calculation protocol within the CS-Rosetta Toolbox 3.0 (*50*).

Selecting the $CytR^{N}$ segment for structure calculation. Order parameters derived from excited state chemical shifts (Fig. 2E) indicate that $S^2$ values are smaller than 0.5 at the N-terminus (M1-M12) and C-terminus (P57-E66). The trends in order parameters agree with chemical shift perturbations (Fig. 2D), which indicate that residues before T11 and beyond Q56 do not change significantly in chemical shift when the native state transitions to the excited conformation. Accordingly, we calculated structures for two segments of $CytR^{N}$, 11-53 and 11-57. Since the structures of the two segments are virtually identical (Fig. S8A) and the region between 54-57 does not participate in secondary structure or tertiary interactions, all subsequent descriptions below pertain to the structure calculation protocol for $CytR^{N}$(11-53).

Fragment picking. As the first step, a fragment library containing 200 3-amino acid fragments and 200 9-residue fragments was constructed for each residue position of $CytR^{N}$(11-53) using the pick_fragments module(*50*). 34 $^{15}N$, 33 $^{1}H^{N}$, 25 $^{13}C\alpha$ and 27 $^{13}C'$ chemical shifts were used as inputs at this stage. Fragments are chosen from a database containing segments of X-ray structures using the similarity of the secondary structure predicted by TALOS-N (*73*) for the target sequence with the X-ray structure as the primary guide. The selected fragments are subsequently scored based on how well their sequence, secondary structure and chemical shifts (predicted by Sparta+)(*51*) match with the target. Homologous structures were eliminated from the database during fragment picking using the -nohom flag in order to avoid biasing the structure calculation.

Fragment assembly. The fragment library was then used along with 119 chemical shifts and 65 RDCs as inputs for the fragment assembly with the 'abrelax' tool(*50*). Structure calculation with Abrelax consists of two stages: in the first stage (Abinitio), a coarse-grained conformational search is performed to obtain low-resolution structural models that are scored using a 'centroid' scoring function; in the next stage (Relax), the structure undergoes all-atom refinement in the Rosetta force field. The CS-Rosetta scoring function was modified to incorporate an energy term arising from deviations of predicted RDCs from experimental values. The weighting factor for this deviation was adjusted so that the width of the resulting distributions of energy contributions from chemical shifts and RDCs are approximately equal (*74*). The RDC weighting factor was the same for both the Abinitio and the Relax stages of structure calculation.

Analysis of structures. 10000 structures of CytR$^N$(11-53) were calculated using the above protocol. The structures were then rescored according to the deviations of their Sparta+-predicted chemical shifts from the experimental values using Eq. 1 of Shen *et al.* (*49*). The energy vs RMSD (to the lowest energy structure) shows a well-defined convergence funnel (Figure S8B) The 10 structures (*74*) with the lowest scores were used as descriptors of the excited state ensemble. The quality of the structures was checked with the PSVS Validation Suite(*75*).

In order to determine how well the excited state structure agrees with input experimental data, we first back-predicted the $^{15}$N, $^{1}$H$^N$, $^{13}$C$\alpha$ and $^{13}$C' chemical shifts for the lowest energy conformer from CS-Rosetta using Sparta+. The shifts agree well with the experimental values (Fig. S9) and the RMSDs for all nuclei ($^{15}$N: 1.95 ppm, $^{1}$H$^N$: 0.38 ppm, $^{13}$C': 1.09 ppm, $^{13}$C$\alpha$: 0.58 ppm) are within the performance of the Sparta+ neural network (*51*) ($^{15}$N: 2.45 ppm, $^{1}$H$^N$: 0.49 ppm, $^{13}$C': 1.09 ppm, $^{13}$C$\alpha$: 0.94 ppm).

We next used PALES (*76*) to predict the RDCs for the excited state by fitting the measured RDCs to the final excited state structure and using the optimized alignment tensor to evaluate the RDCs. The predicted RDCs agree very well with the experimental RDCs (Fig. S10), with an RMSD of 0.87 Hz for PAG and 1.26 Hz for $C_8E_5$/octanol. The Cornilescu quality factor (Q) values (*77*) for the fits are 0.124 and 0.091 for PAG and $C_8E_5$/octanol respectively, demonstrating that there is good agreement between the experimental RDCs and the RDCs calculated from the excited state structure.

Flux calculations. The flux of molecules reaching the specific DNA-CytR$^N$ complex through the D→B and E→B pathways was calculated from the rate constants ($k_{ex,DB}$, $k_{ex,EB}$) and populations ($p_E$, $p_B$) obtained by fitting D- and DRD-CEST data as described in Materials and Methods, along with the total protein ($P_T$) and DNA concentration ($D_T$) in Sample 4. The rate constant $k_{ex,EB}$ is related to the association rate constant between the excited state and DNA, $k_{on,EB}$ as:

$$k_{on,EB} = \frac{k_{EB}}{[DNA]} = \left(\frac{k_{ex,EB}}{[DNA]}\right)\left(\frac{p_B}{p_B + p_E}\right)$$

Here,

$$k_{ex,EB} = k_{EB} + k_{BE} = k_{on,EB}[DNA] + k_{BE}$$

Similarly,

$$k_{on,DB} = \frac{k_{DB}}{[DNA]} = \left(\frac{k_{ex,DB}}{[DNA]}\right)\left(\frac{p_B}{p_B + p_D}\right)$$

Here, $[i]$ is the equilibrium concentration of species $i$. From the mass balance equations,

$$[DNA] = D_T - [B] = D_T - P_T p_B$$

The flux along the E→B and D→B pathways are given by:

$$\phi_{EB} = k_{on,EB}[E][DNA]$$

and

$$\phi_{DB} = k_{on,DB}[D][DNA]$$

The ratio of the flux along the D→B and E→B pathways (Φ) is then given by:

$$\Phi = \frac{\phi_{EB}}{\phi_{DB}} = \frac{k_{on,EB}[E][DNA]}{k_{on,DB}[D][DNA]} = \left(\frac{k_{ex,EB}}{k_{ex,DB}}\right)\left(\frac{p_E}{p_D}\right)\left(\frac{p_D + p_B}{p_E + p_B}\right)$$

We calculated the flux ratio F in four different ways.

1. We first calculated Φ for the rate constant values obtained directly from the fit. Using the values $k_{ex,DB}$ = 0.00015 $s^{-1}$, $k_{ex,EB}$ = 194 $s^{-1}$, $p_E$ = 0.16, $p_B$ = 0.02 and $p_D = 1 - (p_E + p_B) = 0.82$, the ratio of flux along the E→B to the D→B pathways comes out to be 1.2 x $10^6$.
2. Since $p_E$ is slightly different from the value obtained in the absence of DNA because it was allowed to vary during the triangular fitting routine, we next calculated Φ using the $p_E$ value determined from $^{15}$N CEST experiments on DNA-free wt CytR (Sample 1, $p_E$ = 0.087). $p_E$ is a fractional ratio such that the sum of the populations of all conformations in the system is 1. Therefore, from the data on Sample 1,

$$\frac{p_E}{p_D} = \frac{0.087}{1 - 0.087} = 0.095$$

   Using this ratio for the data on Sample 4, where three conformations D, E and B coexist in equilibrium, we get $p_E$ = 0.085, $p_B$ = 0.02 and $p_D = 1 - (p_E + p_B) = 0.89$. The flux ratio Φ from these populations, as well as the rate constants from above ($k_{ex,DB}$ =

0.00015 $s^{-1}$, $k_{ex,EB}$ = 194 $s^{-1}$), is evaluated to be 1.1 x $10^6$, which is very similar to the value calculated in method 1 above.

The error in $k_{ex,DB}$ (0.3 $s^{-1}$) is much larger than the value itself (0.00015 $s^{-1}$), since the data-driven measure of $k_{ex,DB}$ is very close to 0. In the next two methods, we used a 3σ value as an estimate of the upper bound of $k_{ex,DB}$, where σ (=0.3 $s^{-1}$) is the error in $k_{ex,DB}$.

3. Using $k_{ex,DB}$=3σ=0.9 $s^{-1}$ and the other parameters from method 1 ($k_{ex,EB}$ = 194 $s^{-1}$, $p_E$ = 0.16, $p_B$ = 0.02 and $p_D = 1 - (p_E + p_B) = 0.82$), the ratio of flux along the E→B to the D→B pathways is calculated to be 2.0 x $10^2$.
4. Using $k_{ex,DB}$=3σ=0.9 $s^{-1}$ and the other parameters from method 2 ($k_{ex,EB}$ = 194 $s^{-1}$, $p_E$ = 0.085, $p_B$ = 0.02 and $p_D = 1 - (p_E + p_B) = 0.89$), the ratio of flux along the E→B to the D→B pathways is evaluated to be 1.8 x $10^2$.

Therefore, the limits of Φ imposed by the D- and DRD-CEST data on D34 in conjunction with the triangular model are between 1.8 x $10^2$ and 1.2 x $10^6$.

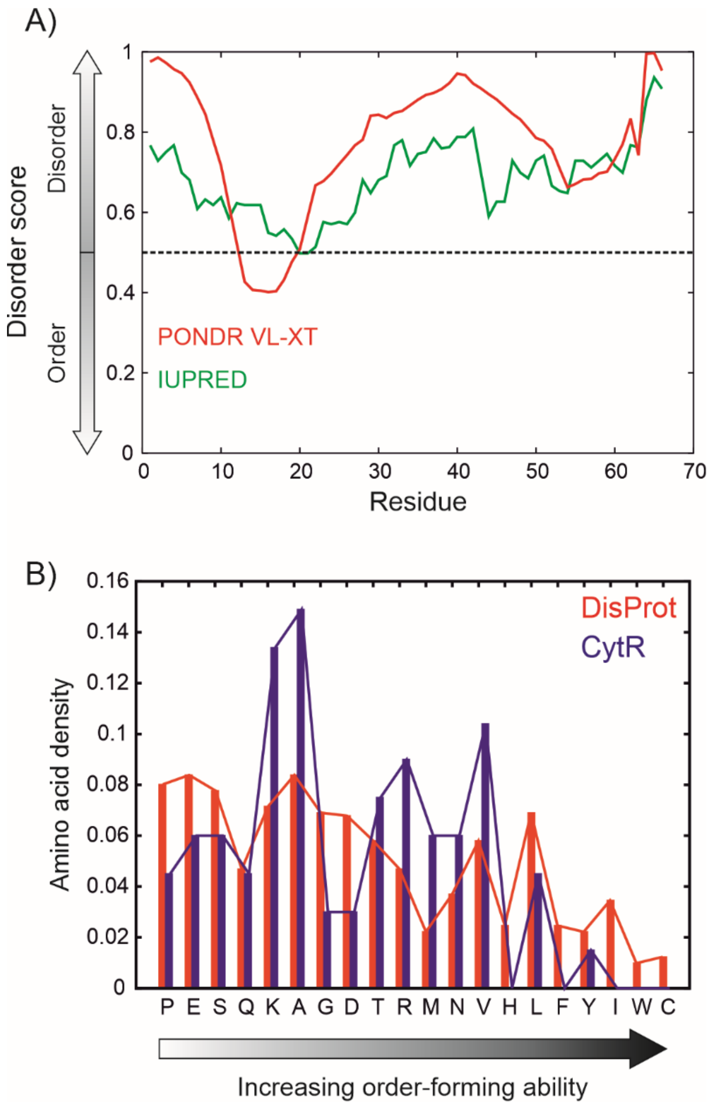


**Fig. S1.**

CytR$^{N}$ is predicted to be intrinsically disordered. A) Residue-specific disorder scores calculated using the PONDR VL-XT (*29, 30*) algorithm (red) and IUPRED (*31*) (green). B) Bar plot of the number of each of the 20 amino acids in CytR$^{N}$ (blue), arranged according to their increasing order-forming tendency. (Red) The frequency of each amino acid in the DisProt database (*78*) of intrinsically disordered proteins.

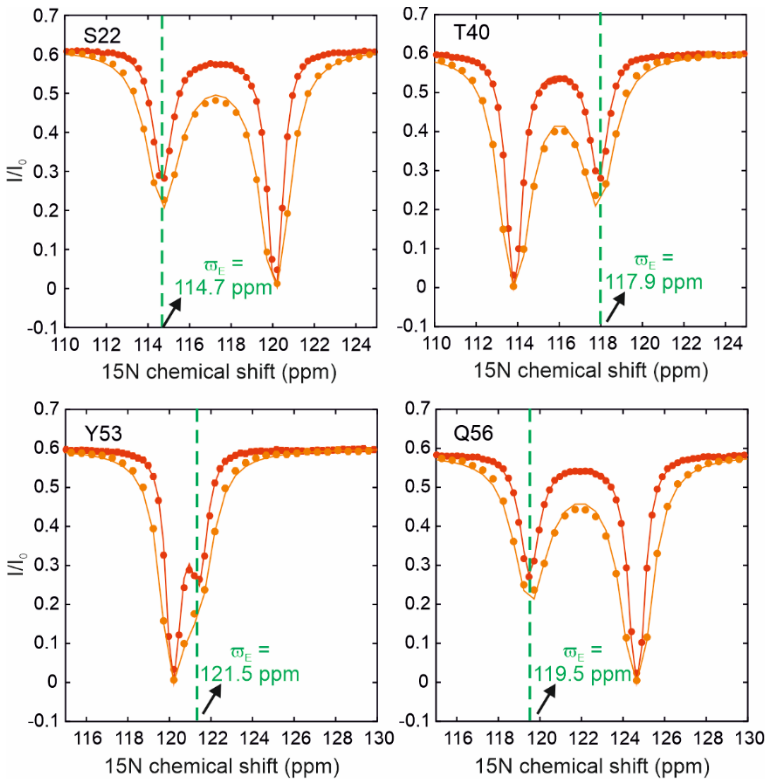


**Fig. S2.**

$^{15}$N CEST profiles of wt CytR$^{N}$ reveal the presence of a conformationally excited state. $^{15}$N CEST profiles of S22, T40, Y53 and Q56 acquired at 14.7 Hz (red) and 28.6 Hz (orange). Solid lines are global fits of the CEST data to the Bloch-McConnell equations for two-site exchange, while the dashed line shows the chemical shift position of the excited state.

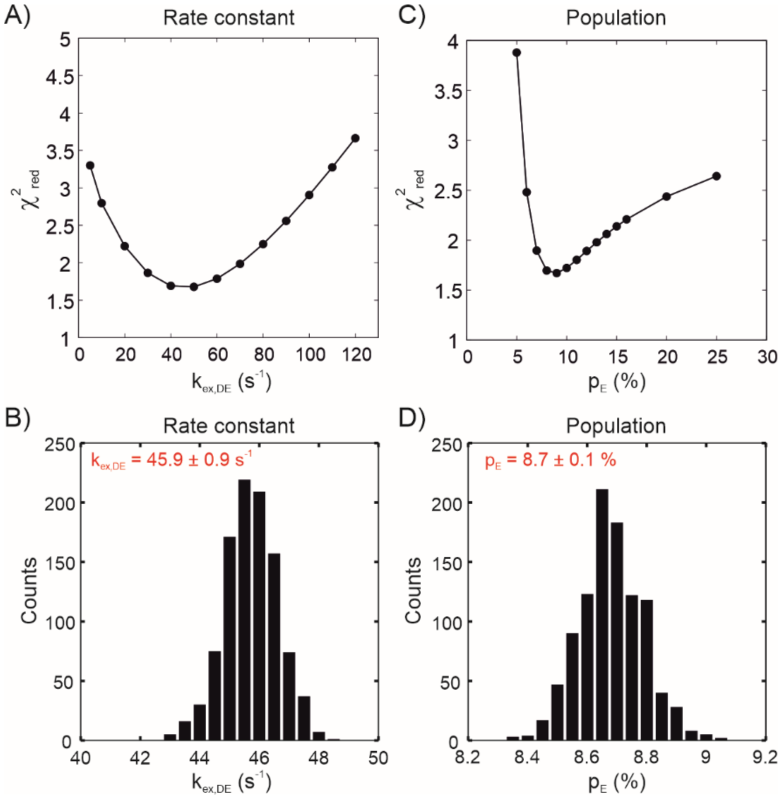


**Fig. S3.**
$\chi^2_{red}$ surfaces (A,C) and Monte Carlo distributions (B,D) for the exchange rate constant between the disordered and excited state ($k_{ex,DE}$; A,B) and the population of the excited state ($p_E$; C,D).

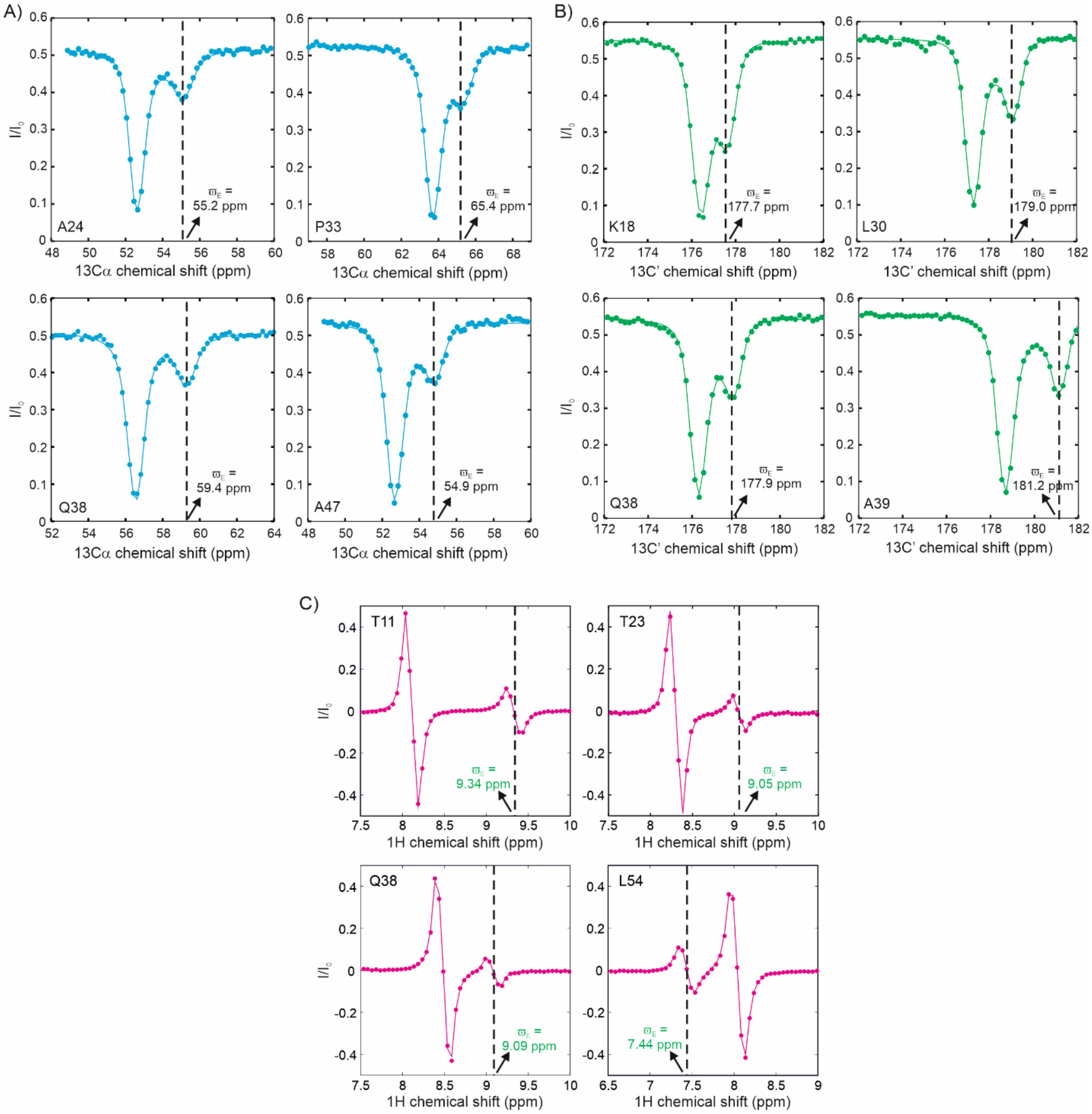


**Fig. S4.**

A) $^{13}$Cα, B) $^{13}$C' and C) $^{1}$H$^{N}$ CEST profiles showing the exchange between the disordered native ensemble and the excited state of CytR$^{N}$. The dashed line shows the chemical shift position of the excited state.

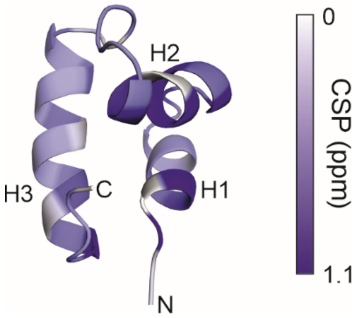


**Fig. S5.**
Cartoon representation of the three-helix bundle structure of DNA-bound $CytR^N$ (PDB ID: 2LCV (*33*)). The backbone has been coloured from white to purple according to the magnitude of the chemical shift perturbation (CSP) of each residue between the disordered and excited state. The color bar mapping the colour with the CSP value is shown alongside.

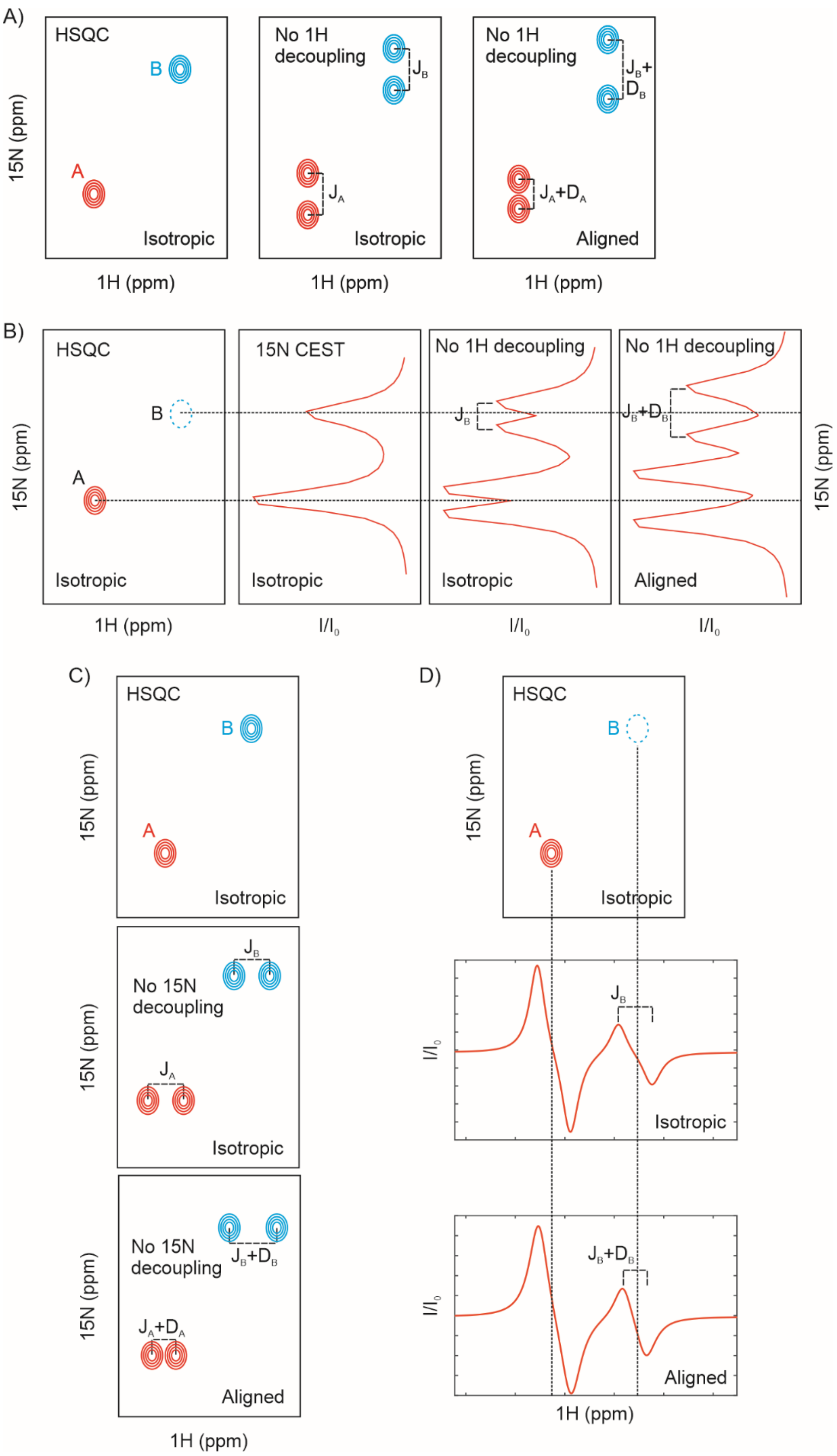


**Fig. S6.**

Schematic of how excited state RDC measurements were made using $^{15}$N and $^{1}$H$^{N}$ CEST profiles. A,C) $^{1}$H$^{N}$-$^{15}$N RDCs for a protein exchanging between two visible conformations A and B can be measured by removing either $^1H^N$ decoupling during $t_1$ (A) or $^{15}N$ decoupling during $t_2$ from a heteronuclear correlation NMR experiment. The splitting between the doublets in an isotropic sample (middle panel) gives $^1J_{NH}$, while the splitting in an aligned sample gives $^1J_{NH}$+RDC (5A, right panel; 5C, bottom panel), with the difference corresponding to the residue-specific RDC value. (B,D) $^{15}N$ and $^1H^N$ CEST profiles can be used to visualize the excited state B which is not observable in heteronuclear correlation spectra and also to determine the RDC values for both states A and B. In the $^{15}N$ CEST experiment (panel B), the major state A and the minor state B both give rise to dips in intensity in an isotropic sample. If the $^1H$ decoupling during the exchange duration is removed, each dip splits into a doublet separated by $^1J_{NH}$ in the isotropic sample and $^1J_{NH}$+RDC in the aligned sample. In the $^1H^N$ CEST experiment the CEST profile is already a difference of two Lorentzians, one corresponding to the TROSY component ($H_zN_\beta$) and the other to the anti-TROSY component ($H_zN_\alpha$). Therefore, the lineshape contains information about $^1J_{NH}$ in the isotropic sample and $^1J_{NH}$+RDC in the aligned sample.

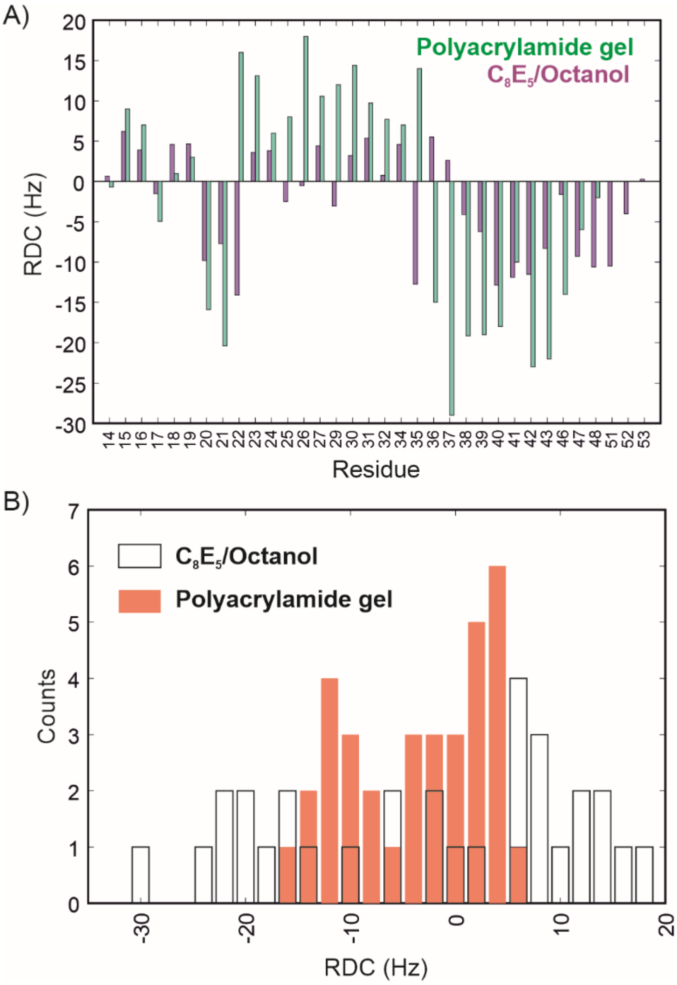


**Fig. S7.**

A) Bar plot of experimentally measured excited state RDCs (aligned – isotropic) in PAG (green) and $C_8E_5$/octanol (purple). B) Histogram of experimentally measured RDCs in PAG (filled orange boxes) and $C_8E_5$/octanol (open boxes).

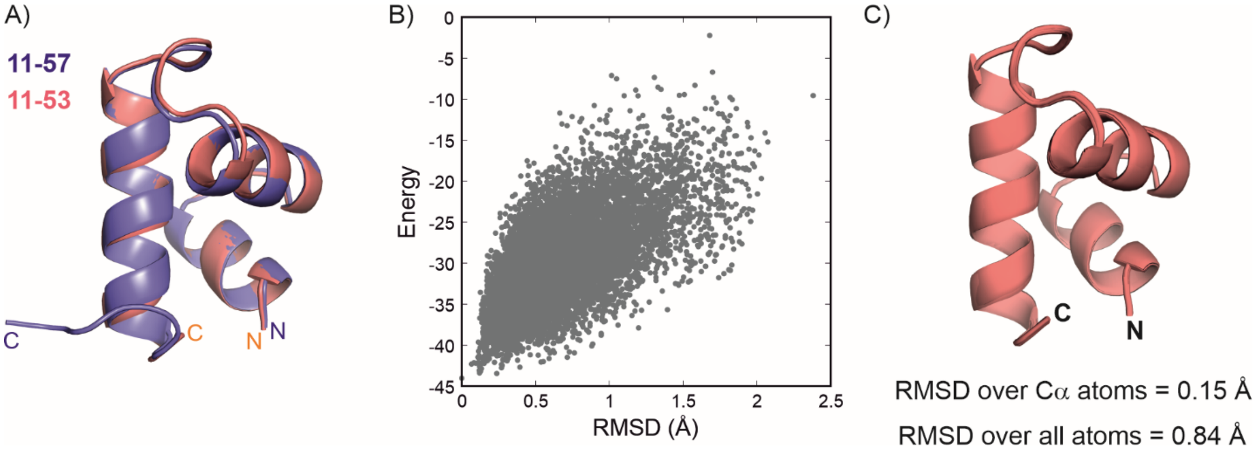


**Fig. S8.**

A) Comparison of the CS-Rosetta structures of CytR$^N$(11-53) (red) and CytR$^N$(11-57) (blue), showing that the three-helix bundle topology is identical in both structures and that the residues from 54-57 do not adopt any specific secondary structure. B) Plot of the energies of the 10000 conformations sampled during the CS-Rosetta structure calculation against the RMSD of each structure with the lowest energy conformation. B) Superposition of the 10 lowest energy conformations generated by CS-Rosetta.

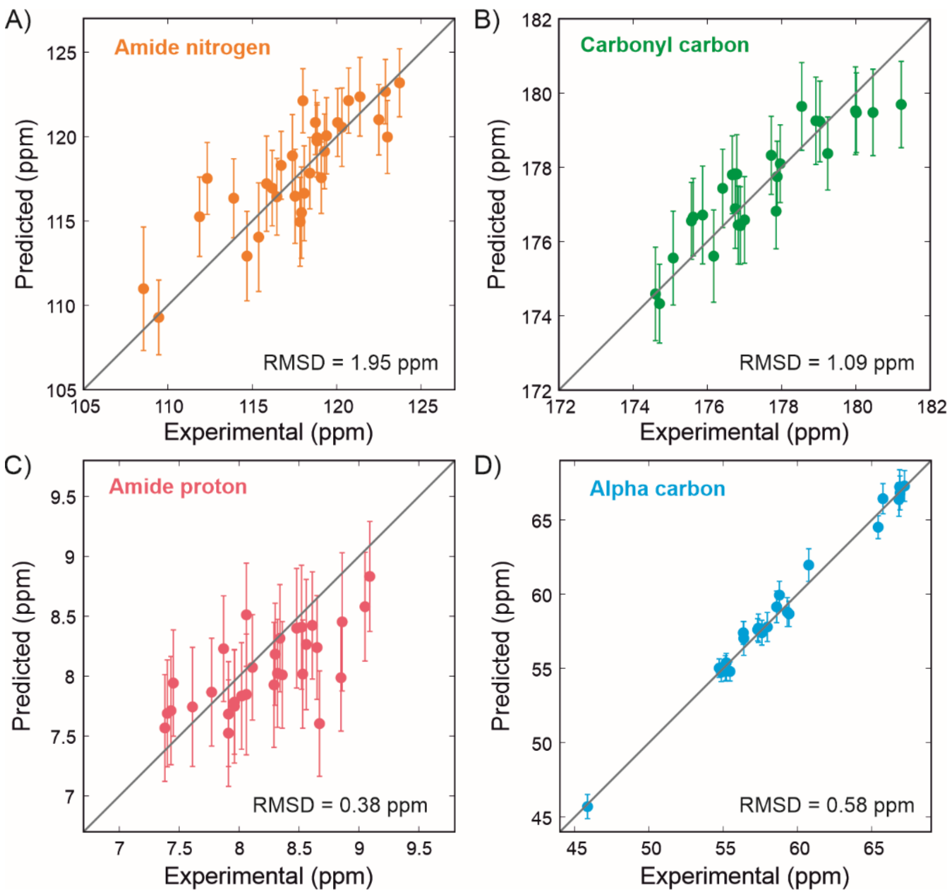


**Fig. S9.**

A comparison of the experimental $^{15}N$ (A), $^{13}C'$ (B), $^{1}H^{N}$ (C) and $^{13}C\alpha$ (D) chemical shifts (x-axis) with the shifts predicted by Sparta+ for the lowest energy conformer from CS-Rosetta (y-axis). The RMSD values for each correlation are indicated on the plot in ppm. The errors in the predicted chemical shifts are obtained directly from Sparta+.

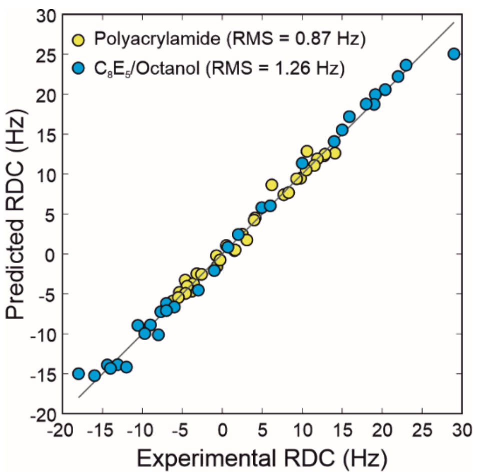


**Fig. S10.**

A comparison of the experimental RDCs (x-axis) in PAG (yellow) and $C_8E_5$/octanol (blue) with the RDCs predicted for the lowest energy CS-Rosetta conformer by PALES (see Materials and Methods). The RMSD values for each correlation are indicated on the plot in Hz.

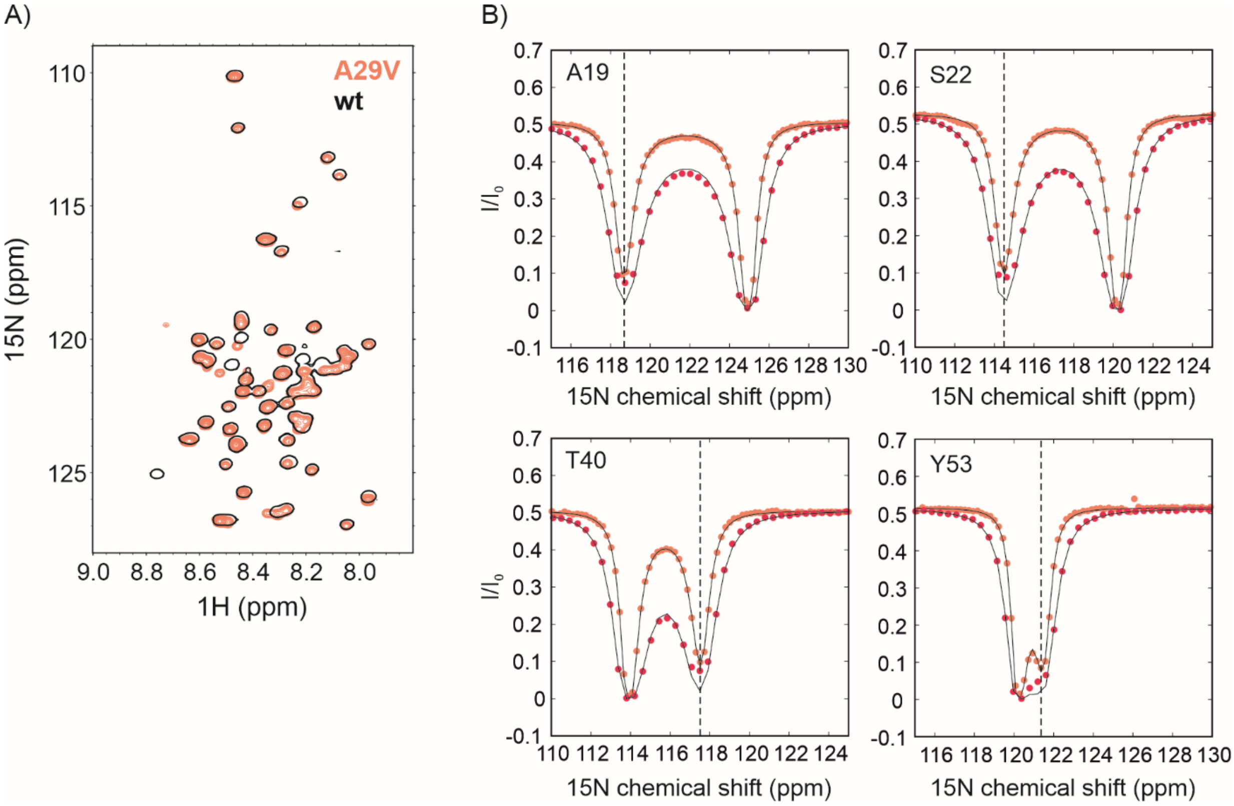


**Fig. S11.**

A) $^{1}$H-$^{15}$N HSQC spectrum of A29V CytR$^{N}$ (orange) overlaid with the spectrum of wt CytR$^{N}$ (black). B) $^{15}$N CEST profiles of A29V CytR$^{N}$ acquired at 12.4 (orange) and 23.9 Hz (red). The black dashed line indicates the position of the excited state.

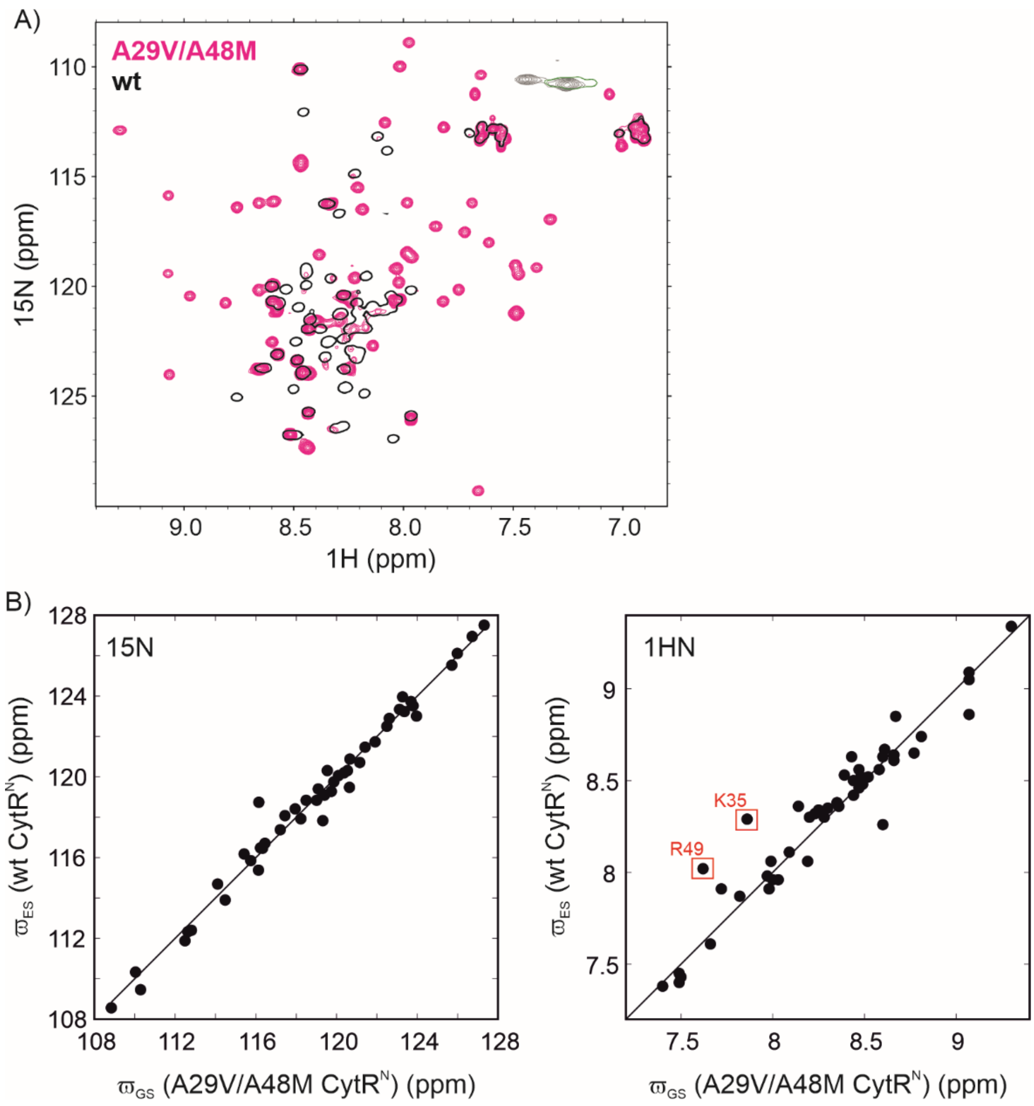


**Fig. S12.**

A) $^{1}$H-$^{15}$N HSQC spectrum of A29V/A48M CytR$^{N}$ (magenta) overlaid with the spectrum of wt CytR$^{N}$ (black). The relative populations of the excited and ground states for A29V/A48M CytR$^{N}$ were obtained directly from the $^{15}$N-$^{1}$H HSQC spectrum, since resonances from both states were visible. Six isolated resonances for which reliable assignments could be obtained for both the excited and disordered conformations were used for analysis. The volumes of these peaks were used to determine the populations of the excited and disordered states. B) Correlations between the $^{15}$N (left) and $^{1}$H$^{N}$ (right) chemical shifts of the folded ground state of A29V/A48M CytR$^{N}$ (x-axis) and the excited state of wt CytR$^{N}$ (y-axis). The solid line is a graph of the y=x function. R49 is adjacent to the site of A48M mutation, which likely results in the 0.4 ppm difference between the $^{1}$H$^{N}$ chemical shifts of the two states. K35 forms a hydrogen bond with N32 in excited state of wt CytR$^{N}$. The upfield shift of K35 $^{1}$H$^{N}$ chemical shift by 0.4 ppm compared to wt $CytR^N$ suggests that this hydrogen bond is likely weakened or broken by the A29V mutation at the end of helix H2.

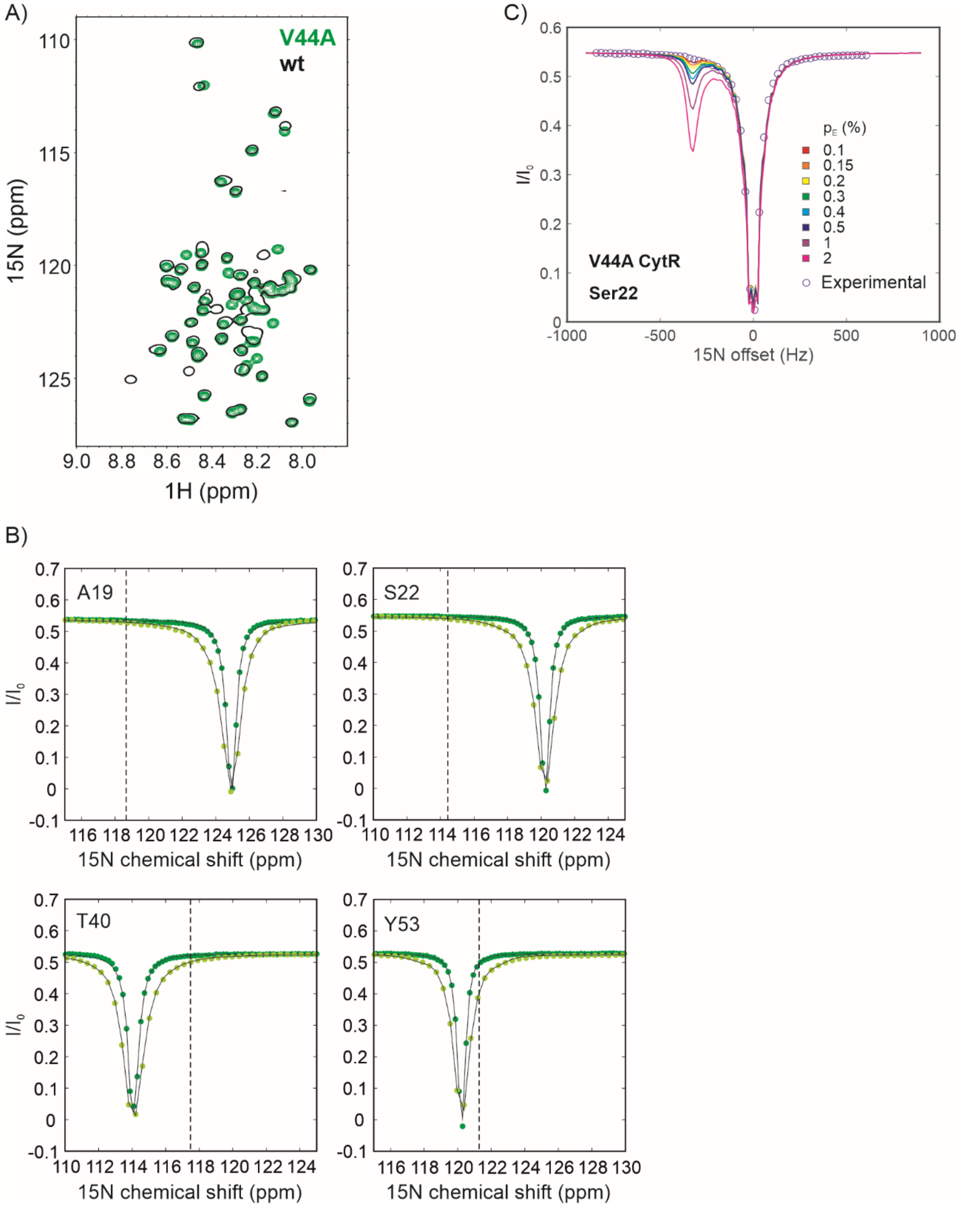


**Fig. S13.**

A) $^{1}$H-$^{15}$N HSQC spectrum of V44A CytR$^{N}$ (green) overlaid with the spectrum of wt CytR$^{N}$ (black). B) $^{15}$N CEST profiles of V44A CytR$^{N}$ acquired at 14.0 (dark green) and 26.5 Hz (light green). The black dashed line indicates the position of the excited state. The minor dip at this position is absent in the CEST profiles of V44A CytR$^{N}$, confirming that the excited state is destabilized in this mutant. C) Obtaining the upper limit for the population of the excited state for V44A CytR$^N$. Open circles indicate experimentally acquired $^{15}N$ CEST data ($B_1$ field of 26.5 Hz), while coloured lines are simulated CEST profiles for various populations ranging from 0.1 to 2 %. From the size of the minor dip, the upper limit of the population is estimated as 0.15 %, since minor dips for populations larger than 0.15 % are expected to be visible in CEST profiles of V44A CytR$^N$.

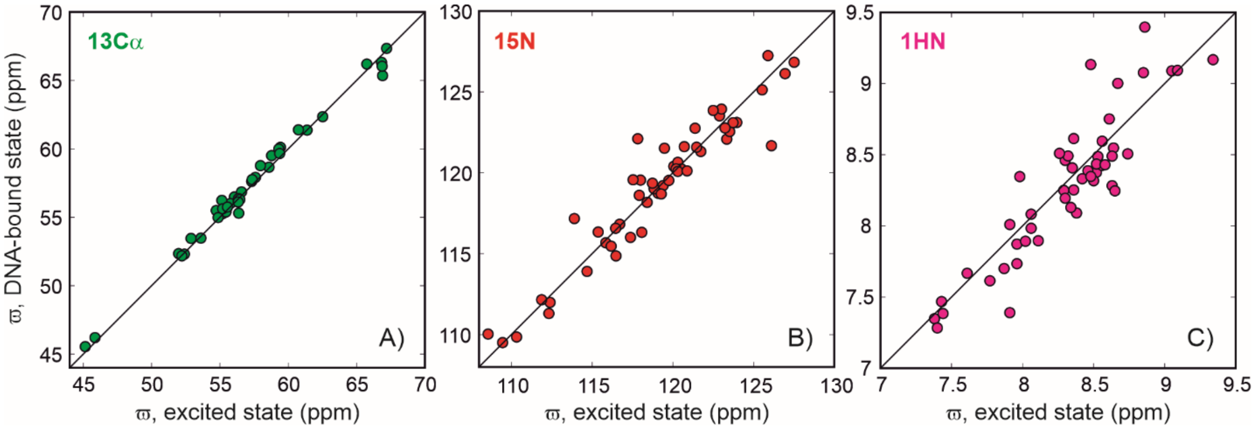


**Fig. S14.**

A comparison of the $^{13}C\alpha$ (A), $^{15}N$ (B) and $^{1}H^{N}$ (C) chemical shifts of the excited state (x-axis) and the DNA-bound state (y-axis) of CytR$^{N}$. Excited state chemical shifts were obtained from CEST profiles, while the shifts of the DNA-bound state are from BMRB ID 17419.

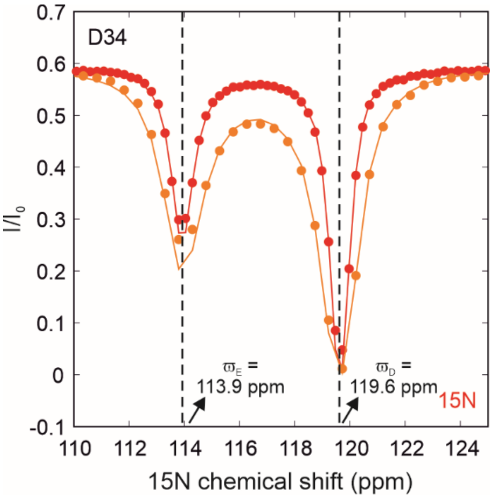


**Fig. S15.**
$^{15}$N CEST profiles of D34 acquired in absence of DNA at 14.7 (red) and 28.6 Hz (orange). The dashed black lines indicate the positions of the disordered and excited states at 119.6 and 113.9 ppm respectively.

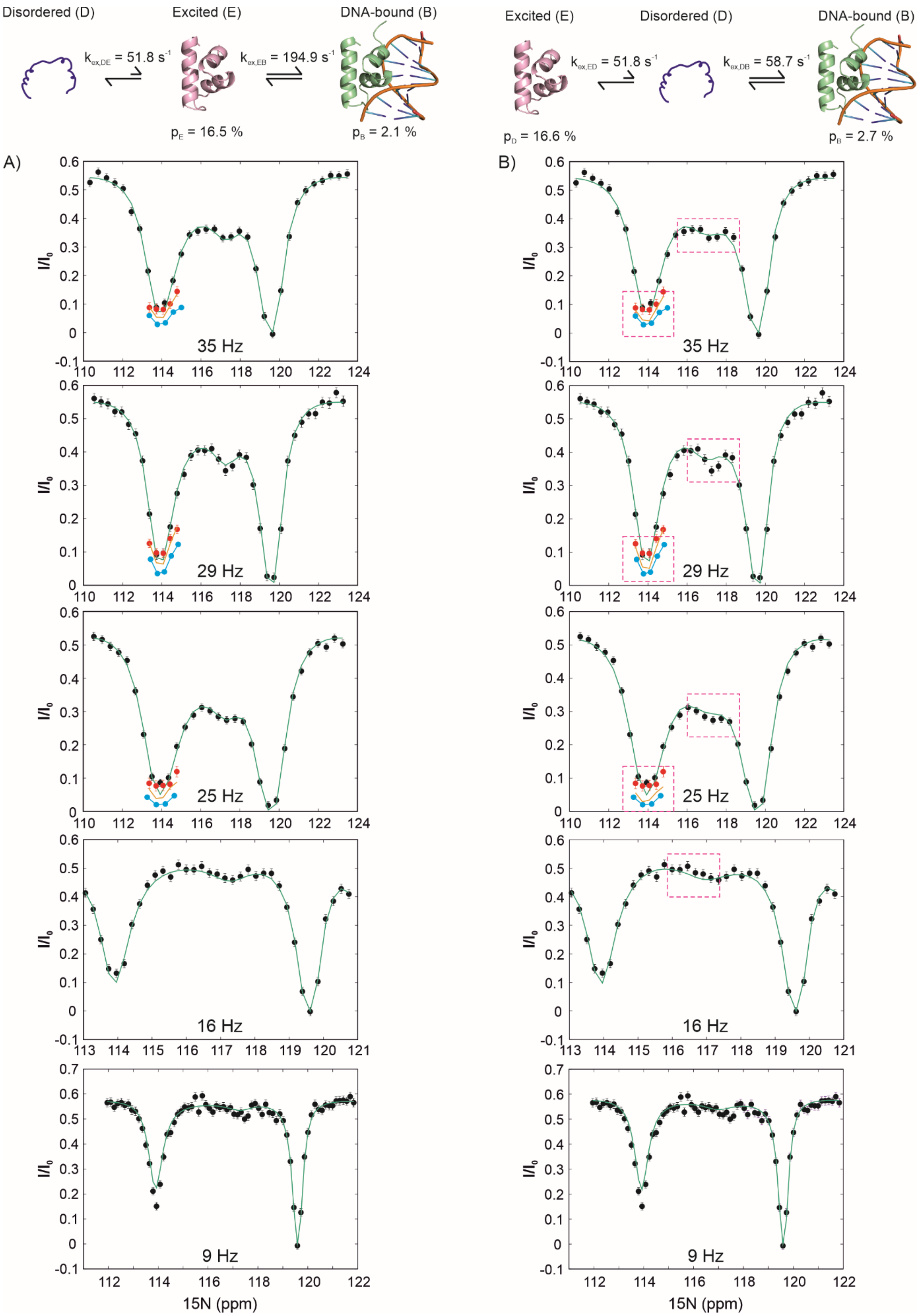
Disordered (D)
Excited (E)
DNA-bound (B)
$k_{ex,DE}$ = 51.8 $s^{-1}$
$k_{ex,EB}$ = 194.9 $s^{-1}$
$p_E$ = 16.5 %
$p_B$ = 2.1 %
Excited (E)
Disordered (D)
DNA-bound (B)
$k_{ex,ED}$ = 51.8 $s^{-1}$
$k_{ex,DB}$ = 58.7 $s^{-1}$
$p_D$ = 16.6 %
$p_B$ = 2.7 %
A)
B)
$I/I_0$
35 Hz
29 Hz
25 Hz
16 Hz
9 Hz
15N (ppm)

**Fig. S16.**

Fits of D- and DRD-CEST data of D34 to the D↔E↔B (A) and E↔D↔B (B) models. Black and red filled circles are D- and DRD-CEST data respectively, acquired at the $B_1$ fields indicated in each plot. The data points are the same between corresponding sub-panels of panels A and B. Green and orange lines are the best global fits 5 $B_1$ field D-CEST and 3 $B_1$ field DRD-CEST data to the D↔E↔B model (A) and E↔D↔B model (B) for DNA binding. Cyan circles are simulated DRD-CEST data assuming the correct exchange mechanism is the E↔D↔B model. Simulations were done using the best-fit parameters obtained by fitting 5 D-CEST and 3 DRD-CEST globally to the E↔D↔B model ($k_{ex,DE}$ = 51.8 $s^{-1}$, $k_{ex,DB}$ = 58.7 $s^{-1}$, $p_E$ = 16.6 % and $p_B$ = 2.7 %). The significant difference between the experimental DRD-CEST data (orange) and the simulated data for the E↔D↔B model (cyan) confirms that the correct mechanism of exchange is the D↔E↔B model. Magenta boxes in panel B highlight systematic deviations between the CEST data and the fits for the E↔D↔B model that are not present in the corresponding fit to the D↔E↔B model. In panel B, the fits systematically underestimate the size of the minor dip at the chemical shift of the DNA-bound state in order to accommodate the expected DRD-CEST data, so that the fits to the DRD-CEST (orange lines) are also systematically deeper than the experimental data (red circles) and closer to the simulated datapoints (cyan).

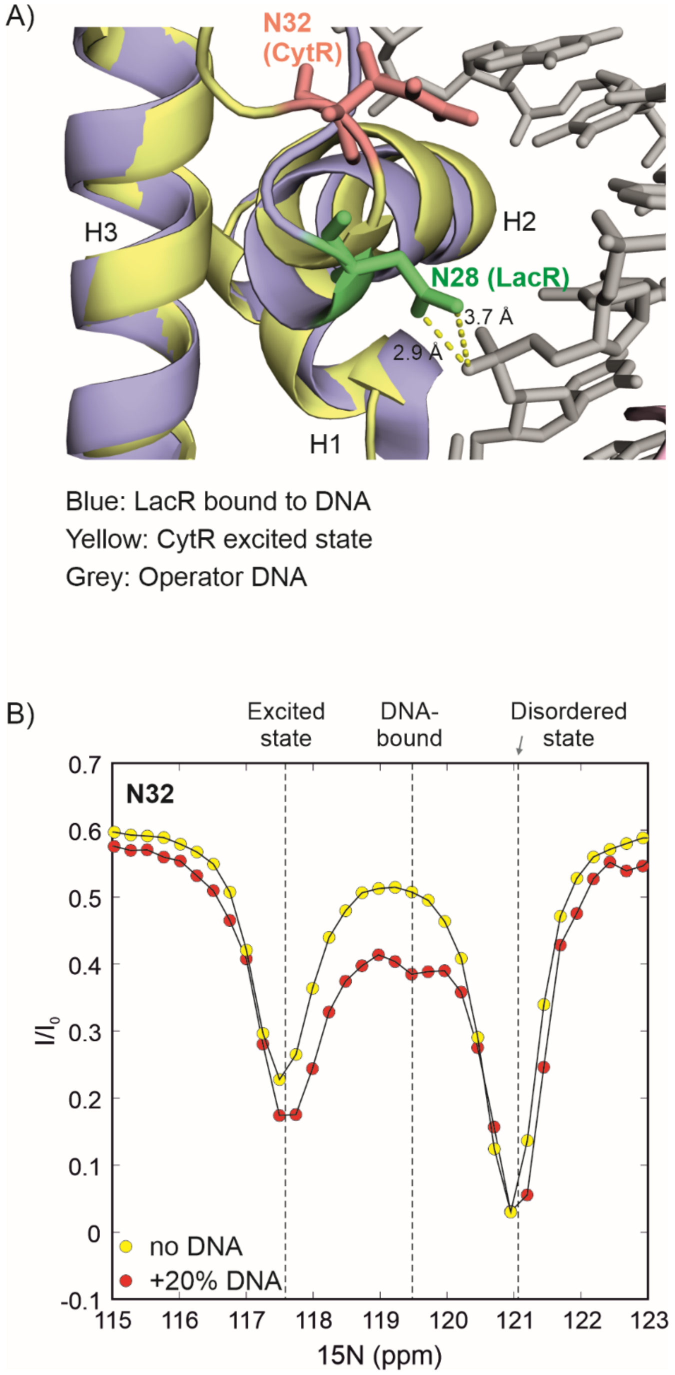


**Fig. S17.**

A) Overlay of DNA-bound LacR (blue, PDB ID 1EFA (*59*)) and the excited state of CytR$^{N}$ (yellow). The DNA molecule is shown as grey sticks. Helix H2 is slightly shorter in the excited state of CytR$^{N}$. The sidechain of N25 in LacR forms hydrogen bonds with the DNA phosphodiester backbone (*59, 60*), while the corresponding residue in CytR$^{N}$ (N32) is further away and oriented differently, suggesting that this residue reorganizes itself after the folded state of CytR$^{N}$ binds DNA. In agreement with this, $^{15}N$ CEST profiles of N32 in the DNA-bound form (B, red) show that N32 has a significantly different chemical shift in the DNA-bound form compared to the excited state.

**Table S1.**
List of wt and mutant samples used for NMR experiments.

| Sample number | Variant | Isotope labelling | Protein conc. (μM) | DNA conc. (μM) | Experiments carried out |
|---|---|---|---|---|---|
| 1 | Wt | $^{13}C$, $^{15}N$ | 614 | - | $^{15}N$ CEST and backbone assignment experiments |
| 2 | Wt | $^{13}C$, $^{15}N$ | 713 | - | $^{13}C\alpha$ CEST, $^{13}C\alpha$ Gly CEST, $^{1}H^{N}$ CEST, $^{13}C'$ CEST |
| 3 | Wt | $^{15}N$ | 648 | 150 | DNA titration |
| 4 | Wt | $^{15}N$ | 628 | 150 | $^{15}N$ D-CEST, $^{15}N$ DRD-CEST |
| 5 | Wt | $^{15}N$ | 1060 | - | Isotropic sample for PAG |
| 6 | Wt | $^{15}N$ | 980 | - | RDC (PAG) |
| 7 | Wt | $^{15}N$ | 793 | - | Isotropic sample for $C_8E_5$/octanol |
| 8 | Wt | $^{15}N$ | 806 | - | RDC $C_8E_5$/octanol |
| 10 | A29V | $^{15}N$ | 1800 | - | $^{15}N$ CEST |
| 11 | V44A | $^{15}N$ | 1300 | - | $^{15}N$ CEST |
| 12 | A29V/A48M | $^{13}C$, $^{15}N$ | 843 | - | $^{15}N$ CEST and backbone assignment experiments |

**Table S2.**

Parameters used for recording multinuclear CEST experiments on wt and mutant CytR$^N$.

| Sample | Experiment | $B_1$ (Hz) | $T_{ex}$ (ms) | sweep (Hz) | No. of planes | Offset spacing (Hz) | $t_1$ (ms) | $t_2$ (ms) |
|---|---|---|---|---|---|---|---|---|
| CytR$^N$ wt | $^{15}$N CEST | 14.7 | 300 | -720 to 720 | 98 | 15 | 64 | 40 |
| | $^{15}$N CEST | 28.6 | 300 | -720 to 720 | 50 | 30 | 64 | 40 |
| | $^{13}$C' CEST | 25 | 200 | -900 to 900 | 62 | 30 | 64 | 38 |
| | $^{1}$H$^N$ CEST | 25 | 300 | -1500 to 1500 | 102 | 30 | 64 | 33 |
| | $^{13}$Cα CEST | 25 | 250 | -1500 to 1500 | 102 | 30 | 64 | 31 |
| | $^{13}$Cα Gly CEST | 25 | 250 | -2220 to 1620 | 22 | 30 | 64 | 31 |
| V44A CytR$^N$ | $^{15}$N CEST | 14.0 | 400 | -720 to 723 | 113 | 13 | 64 | 42 |
| | $^{15}$N CEST | 26.5 | 400 | -720 to 730 | 60 | 25 | 64 | 42 |
| A29V CytR$^N$ | $^{15}$N CEST | 12.4 | 400 | -720 to 723 | 113 | 13 | 64 | 42 |
| | $^{15}$N CEST | 23.9 | 400 | -720 to 730 | 60 | 25 | 64 | 42 |

**Table S3.**
List of chemical shifts[1] of the wt $CytR^N$ excited state.

| Res no. | $\varpi_E$ ($^{15}N$) | $\varpi_E$ ($^{1}H^N$) | $\varpi_E$ ($^{13}C\alpha$) | $\varpi_E$ ($^{13}C'$) |
|---|---|---|---|---|
| 2 | 124.63 | | 56.06 | 176.05 |
| 3 | 126.95 | 8.52 | 51.98 | 177.74 |
| 4 | 121.73 | 8.63 | | |
| 5 | | | | 176.30 |
| 6 | 123.34 | 8.56 | 55.76 | 175.96 |
| 7 | 123.52 | 8.64 | 56.58 | 176.79 |
| 8 | 116.48 | 8.38 | 61.38 | |
| 9 | 127.51 | 8.50 | 52.40 | 178.13 |
| 10 | 123.96 | 8.36 | 52.22 | 178.70 |
| 11 | 112.40 | 9.34 | | |
| 12 | | | | 176.79 |
| 13 | | | 58.78 | |
| 14 | 118.84 | 7.43 | 57.60 | |
| 15 | 119.40 | 7.38 | 65.75 | 176.67 |
| 16 | 122.89 | 8.36 | | |
| 17 | 116.71 | 8.06 | 57.32 | 180.01 |
| 18 | 119.09 | 7.40 | 57.36 | 177.72 |
| 19 | 118.84 | 8.53 | | |
| 20 | 115.38 | 8.06 | | |
| 21 | | | 58.59 | 174.60 |
| 22 | 114.69 | 8.56 | | 176.17 |
| 23 | 115.85 | 9.05 | 66.83 | 176.89 |
| 24 | 123.73 | 8.34 | 55.20 | 180.46 |
| 25 | 118.08 | 7.91 | 66.90 | 175.62 |
| 26 | 121.38 | 7.77 | 67.20 | 176.41 |
| 27 | 111.88 | 8.11 | | |
| 29 | 118.00 | | 54.73 | 178.54 |
| 30 | 112.33 | 7.87 | 56.35 | 179.03 |
| 31 | 116.18 | 8.30 | 56.38 | 176.75 |
| 32 | 117.54 | 8.48 | | |
| 33 | | | 65.43 | 177.00 |
| 34 | 113.90 | 8.52 | 55.14 | 176.83 |
| 35 | 117.38 | 8.29 | | |
| 37 | 117.83 | 8.86 | 57.97 | 175.08 |
| 38 | 123.01 | 9.09 | 59.43 | 177.88 |
| 39 | 120.06 | 8.61 | 55.45 | 181.22 |
| 40 | 117.92 | 7.96 | 66.87 | 175.57 |
| 41 | 122.50 | 8.67 | | |

| | | | | |
|---|---|---|---|---|
| 42 | 116.46 | 8.65 | | 177.85 |
| 43 | | | 59.31 | |
| 44 | 120.31 | 8.32 | | |
| 45 | | | | 177.96 |
| 46 | 119.28 | 7.96 | | |
| 47 | 120.71 | 7.45 | 54.87 | 178.92 |
| 48 | 118.74 | 8.85 | 55.20 | 179.99 |
| 49 | 118.41 | 8.02 | | |
| 50 | | | | 179.24 |
| 51 | 108.56 | 7.91 | 60.75 | 175.87 |
| 52 | 109.46 | 7.61 | 45.86 | 174.71 |
| 53 | 121.47 | 8.35 | 59.37 | 173.98 |
| 54 | 125.89 | 7.44 | | |
| 55 | | | | 176.71 |
| 56 | 119.48 | 8.74 | | |
| 57 | | | | 177.16 |
| 58 | 120.53 | 8.58 | 55.55 | 177.10 |
| 59 | 110.33 | 8.46 | 45.15 | 174.18 |
| 60 | 120.20 | 8.30 | 56.36 | 176.13 |
| 61 | 119.75 | 8.26 | 52.89 | 175.26 |
| 62 | 120.31 | | 62.52 | 176.22 |
| 63 | 125.53 | 8.42 | 56.46 | 176.41 |
| 64 | 123.23 | 8.48 | 56.34 | 175.63 |
| 65 | 120.88 | 8.63 | 53.61 | 174.23 |
| 66 | 126.11 | 7.98 | | |

[1] The average errors in the chemical shifts of the excited state extracted with ChemEx were of the order of 0.01 ppm in $^{15}N$, 0.003 ppm in $^{1}H^{N}$, 0.03 ppm in $^{13}C\alpha$ and 0.09 ppm in $^{13}C'$.

**Table S4.**

Parameters used for recording $^{15}N$ and $^{1}H^{N}$ CEST profiles in isotropic and aligned media.

| Sample No. | Sample No. | Spectrometer (MHz) | Experiment | $B_1$ (Hz) | CEST sweep (Hz) | $T_{ex}$ (ms) | No. of planes |
|---|---|---|---|---|---|---|---|
| 4 | Isotropic (for polyacrylamide) | Agilent 600 MHz | $^{15}N$ CEST (coupled) | 8 | -750 to +750 | 400 | 127 |
| | | | $^{1}H^{N}$ CEST | 15 | -1000 to +1000 | 500 | 102 |
| 5 | Aligned (polyacrylamide stretched gel) | Agilent 600 MHz | $^{15}N$ CEST (coupled) | 8 | -750 to 750 | 400 | 127 |
| | | | $^{1}H^{N}$ CEST | 15 | -1000 to +1000 | 500 | 102 |
| 6 | Isotropic (for $C_8E_5$/octanol) | Bruker 700 MHz | $^{15}N$ CEST (coupled) | 10 | -876 to +876 | 400 | 148 |
| | | | $^{1}H^{N}$ CEST | 15 | 1370 to 3470 | 500 | 107 |
| 7 | Aligned ($C_8E_5$/octanol) | Bruker 700 MHz | $^{15}N$ CEST (coupled) | 10 | -876 to +876 | 400 | 148 |
| | | | $^{1}H^{N}$ CEST | 15 | 1370 to 3470 | 500 | 107 |

**Table S5.**
List of $^{15}N$-$^{1}H$ RDCs of the excited state of CytR$^{N}$ used in CS-Rosetta structure calculations.

| Residue number | Polyacrylamide gel (PAG) | Residue number | $C_8E_5$/n-octanol |
|---|---|---|---|
| T11 | 1.1±1.9 | T11 | 6.0±1.8 |
| D14 | 0.7±3.1 | D14 | -0.7±3.4 |
| V15 | 6.2±1.5 | V15 | 9.0±2.9 |
| A16 | 3.9±1.4 | A16 | 7.0±1.9 |
| L17 | -1.5±0.8 | L17 | -4.9±1.8 |
| K18 | 4.6±1.4 | K18 | 1.0±1.8 |
| A19 | 4.6±1.5 | A19 | 3.0±2.8 |
| K20 | -9.8±2.4 | K20 | -15.9±4.3 |
| V21 | -7.7±1.5 | S22 | -20.4±3.6 |
| S22 | -14.1±0.6 | T23 | 16.0±2.6 |
| T23 | 3.6±1.4 | A24 | 13.1±2.0 |
| A24 | 3.8±3.4 | T25 | 6.0±2.2 |
| T25 | -2.5±1.7 | V26 | 8.0±3.5 |
| V26 | -0.5±1.4 | S27 | 18.0±2.2 |
| S27 | 4.4±1.8 | A29 | 10.6±4.4 |
| A29 | -3.1±3.3 | L30 | 12.0±2.9 |
| L30 | 3.2±0.8 | M31 | 14.4±2.3 |
| M31 | 5.4±2.4 | N32 | 9.7±2.6 |
| N32 | 0.7±0.7 | D34 | 7.7±2.4 |
| D34 | 4.6±2.5 | V36 | 7.0±1.9 |
| K35 | -12.7±1.3 | S37 | 14.0±2.2 |
| V36 | 5.5±1.7 | Q38 | -15.0±3.1 |
| S37 | 2.6±1.3 | A39 | -29.0±2.8 |
| Q38 | -4.1±1.9 | T40 | -19.2±3.1 |
| A39 | -6.2±1.6 | R41 | -19.0±2.9 |
| T40 | -12.9±0.8 | R43 | -18.0±2.1 |
| R41 | -11.9±1.5 | K46 | -10.0±2.3 |
| N42 | -11.5±1.1 | A47 | -23.0±9.9 |
| R43 | -8.3±1.5 | A48 | -22.0±4.5 |
| K46 | -1.6±1.6 | V51 | -14.0±4.4 |
| A47 | -9.3±1.6 | G52 | -6.0±1.9 |
| A48 | -10.6±1.7 | Y53 | -2.0±2.4 |

| | |
|---|---|
| V51 | -10.5±2.1 |
| G52 | -4.0±1.2 |
| Y53 | 0.3±1.2 |

**Table S6.**
Parameters used for recording D- and DRD-CEST on Sample 4.

| | Experiment | $B_1$ (Hz) | $T_{ex}$ (ms) | sweep (Hz) | No. of planes | Offset spacing (Hz) | $sw_{DANTE}$ (Hz) |
|---|---|---|---|---|---|---|---|
| 1 | $^{15}$N D-CEST | 8.8 | 300 | -350 to 350 | 72 | 10 | 700 |
| 2 | $^{15}$N D-CEST | 15.9 | 300 | -272 to 272 | 36 | 16 | 544 |
| 3 | $^{15}$N D- CEST | 24.8 | 300 | -450 to 450 | 38 | 25 | 900 |
| 4 | $^{15}$N D- CEST | 29.0 | 300 | -465 to 465 | 33 | 30 | 930 |
| 5 | $^{15}$N D- CEST | 34.7 | 300 | -450 to 450 | 32 | 30 | 900 |
| 6 | $^{15}$N DRD- CEST | 25.0 | 300 | -350 to -100 | 12 | 25 | 254 |
| 7 | $^{15}$N DRD- CEST | 29.0 | 300 | -350 to -100 | 12 | 25 | 254 |
| 8 | $^{15}$N DRD- CEST | 34.9 | 300 | -350 to -100 | 12 | 25 | 254 |
